# USP7 sustains PAX3::FOXO1 enhancer reprogramming and represents a therapeutic vulnerability in Rhabdomyosarcoma

**DOI:** 10.64898/2026.06.03.724934

**Authors:** Pablo Táboas, Enrique Blanco, Estela Prada, Tobias Faehling, María Sánchez-Jiménez, Carla Rios-Astorch, Merce Baulenas-Farres, Berta Estrada-Bes, Xenia Cuadros-Hernández, Silvia Mateo-Lozano, Soledad Gómez-González, Sara Perez-Jaume, Cinzia Lavarino, Thomas G. P. Grünewald, Florencia Cidre-Aranaz, Jaume Mora, Luciano Di Croce, Sara Sánchez-Molina

## Abstract

Rhabdomyosarcoma (RMS) is the most common soft tissue sarcoma in children and is often associated with dismal outcomes, underscoring the urgent need for new therapeutic strategies. RMS arises from embryonic skeletal muscle precursor cells that fail to complete the myogenic differentiation program. Fusion-positive rhabdomyosarcoma (FP-RMS), defined by the presence of recurrent gene fusions such as *PAX3::FOXO1* or *PAX7::FOXO1*, is associated with the poorest overall survival. The encoded fusion oncoprotein cause epigenetic reprogramming that defines the biology and behavior of FP-RMS. Polycomb repressive complex 1 (PRC1)-mediated chromatin regulation contributes to the control of developmental gene programs. Here, we investigate the dependency of RMS on epigenetic remodeling mediated by the PRC1.1 subunits ubiquitin specific protease 7 (USP7) and really interesting new gene 1B (RING1B). We found that USP7 is overexpressed in RMS samples, and that high expression correlates with poor patient prognosis. USP7 and RING1B bind to H3K27ac-enriched regions and colocalize with PAX3::FOXO1 at active enhancers controlling key tumorigenic genes in FP-RMS. Moreover, both shRNA-mediated depletion and pharmacological inhibition of USP7 downregulate PAX3::FOXO1 enhancer-driven genes, induce skeletal muscle differentiation and significantly inhibit FP-RMS tumor growth *in vivo.* Altogether, our findings identify USP7 as a critical regulator of PAX3::FOXO1-bound enhancers and highlight a novel therapeutic opportunity in RMS based on epigenetic dependencies.

## Introduction

Rhabdomyosarcoma (RMS) is the most common soft-tissue sarcoma in children^1,2^. RMS resembles undifferentiated skeletal muscle cells, consistent with a failure of normal myogenic differentiation program. Despite expressing key myogenic transcription factors (MTFs) such as MYOD1 and MYOG, RMS cells are unable to properly execute the myogenic transcriptional program required for terminal differentiation^3^.

Historically, according to the histological, genetic and clinical characteristics, RMS tumors are classified into two major subtypes: embryonic (ERMS) and alveolar (ARMS)^2^. ERMS is more frequent in young children (under 10 years of age) and is mostly found in head, neck and the genitourinary system. This subtype is characterized by alterations in cell cycle genes and the RAS pathway. In contrast, ARMS is predominantly found in adolescents and young adults, and typically located in the extremities and torso. ARMS is defined by the presence of a chromosomal translocation associated with worse prognosis^4–6^. This structural rearrangement generates a fusion gene, either *PAX3::FOXO1* or *PAX7::FOXO1*. *PAX3::FOXO1* is more common (70% of the cases) and confers an even more aggressive phenotype than *PAX7::FOXO1* ^7^. In order to more precisely define the biology and genomic characteristics of RMS, the current accepted classification relies on the presence (fusion positive, FP-RMS) or absence (fusion negative, FN-RMS) of the fusion gene^2,6^. In FP-RMS, the PAX3::FOXO1 fusion protein functions as a potent transcriptional regulator that drives oncogenic transcriptional programs^2,8^. PAX3::FOXO1 oncogene recruits transcription factors, such as MYOD1, MYOG and MYCN, conforming super-enhancer (SEs) structures marked with H3K27ac that activate gene expression and establish the epigenetic environment conducive to tumorigenesis^8^.

Polycomb Repressive Complexes 1 and 2 (PRC1 and PRC2, respectively) are fundamental epigenetic regulators that control gene expression programs during embryonic development and cell fate decisions, including those governing myogenic differentiation^9–17^. PRC1 plays a central role in maintaining cell identity, primarily by repressing genes involved in development and differentiation. The core PRC1 is composed of a Polycomb group protein (PCGF1-6) and an E3-ubiquitin ligase protein (RING1A or B)^18^, whose catalytic activity deposits the repressive histone mark H2AK119ub^10,18^. Mechanistically, PRC1 has been described to recognize H3K27me3, a mark catalyzed by PRC2, thereby contributing to chromatin compaction and gene silencing during differentiation^18^. More recently, it has been proposed that H2AK119ub can also prime chromatin for H3K27me3 deposition, reinforcing transcriptional repression^19,20^.

The canonical PRC1 complex, containing one variant each of PHC and CBX in addition to the catalytic core, has been primarily linked to maintaining gene repression. However, emerging studies indicate that RING1B-containing PRC1 complexes may also facilitate transcriptional activation. Remarkably, recruitment of RING1B to active enhancers occurs across multiple cancers such as melanoma, leukemia and breast cancer, highlighting an activating role in their oncogenic transcriptional programs^21–23^. In melanoma, PRC1 plays a transcriptional activating function with the ubiquitously transcribed tetratricopeptide repeat, X chromosome (UTX) and the acetyltransferase p300^21^. Previous results from our group have demonstrated that RING1B cooperates at active enhancers with EWSR1::FLI1, the pathognomonic fusion oncogene in Ewing sarcoma (EwS)^24^. RING1B mediates EWSR1::FLI1 recruitment to chromatin, facilitating the expression of key oncogene targets^24^. Similarly, the non-canonical PRC1.1 complex, has been reported to mediate the chromatin binding of SS18::SSX, the oncogenic fusion protein that drives Synovial Sarcoma (SySa)^25–27^.

Besides the catalytic core, the non-canonical PRC1.1 complex contains the specific subunits BCOR, RYBP/YAF2, USP7 and KDM2B, whose CXXC DNA-binding domain recognizes unmethylated CpG islands. USP7 is an integral component of PRC1.1, where its deubiquitinating activity stabilizes several complex subunits, including KDM2B^28–30^. It participates in a variety of biological processes, protecting proteins from degradation, including regulation of the p53-MDM2 axis and promotion of mitosis via CDK1 regulation^31,32^ ^33^. Moreover, USP7 has been shown to regulate the stability of MYC family proteins, highlighting a potential role for USP7 in sustaining MYC-driven tumorigenesis in cancers, such as neuroblastoma^34,35^. In leukemia, USP7 has been reported to regulate gene transcription by maintaining the stability and integrity of PRC1.1 complex^28^. These roles make USP7 a promising target for cancer treatment. Accordingly, numerous preclinical studies have identified different USP7 inhibitors with therapeutic potential^36,37^.

Here, we investigate the contribution of USP7 and RING1B, subunits of the PRC1.1 complex, to RMS progression. We show that RMS tumors express USP7 at high levels, which correlate with worse patient prognosis. We generate the genome-wide maps of USP7 and RING1B occupancy in FP-RMS, revealing that the PRC1.1 complex localizes to PAX3::FOXO1-bound enhancers. Genetic silencing of these PRC1.1 subunits as well as USP7 pharmacological inhibition disrupts the transcriptional program driven by PAX3::FOXO1 oncoprotein in FP-RMS cell lines and a patient-derived tumor cell culture. This reprogramming leads to decreased FP-RMS viability *in vitro* and *in vivo*, highlighting USP7 as a promising therapeutic target in RMS.

## RESULTS

### USP7 is overexpressed in FP-RMS and has prognostic value

PRC1 and PRC2 play a key role in regulating transcriptional programs during myogenic differentiation^13,14,17,38^. However, the role of PRC1.1 in RMS tumorigenesis remains poorly defined. To address this, we analyzed the expression of *USP7* and *RING1B* in publicly available repositories such as Gene Expression Omnibus (GEO) dataset (see Methods for GSE numbers). The aggregated dataset contains 58 RMS patient samples (25 FN-RMS and 33 FP-RMS)^39^, 193 samples from other pediatric sarcomas (EwS, osteosarcoma [OS] and SySa) and 49 normal muscle samples at different stages of differentiation (**Supplementary Table 1**). Interestingly, we observed that the expression of *USP7* (probe 201498_at) is higher in FP-RMS than in differentiated pediatric skeletal muscle (p = 0.0481) or other pediatric sarcomas, such as EwS (p = 0.0196), OS (p < 0.0001) and SySa (p = 0.0073) (**Figure 1A**). Remarkably, no significant differences were observed when comparing the expression of *USP7* in FN-RMS to other sarcomas or healthy muscle samples except for OS (p < 0.0001). The expression of the PRC1 catalytic subunit *RING1B* (probe 25215_at) was higher in FN-RMS than in FP-RMS (p < 0.0001), EwS (p < 0.0001) and OS (p < 0.0001), while no differences were observed with SySa (**Figure 1B**). No significant differences in RING1B expression were detected between FP-RMS, EwS, and OS; however, expression levels were lower in FP-RMS than in SySa (p = 0.0171). Expression data of cell lines obtained from the Cell Line Protein Atlas^40^ did not show significant difference when comparing either FP-RMS or FN-RMS to the other tumor entities although USP7 levels were higher in FP-RMS than in FN-RMS (**Supplementary Figure 1 A-B**). Thus, the higher expression of *USP7* in FP-RMS support further studies to elucidate its role.

**Figure 1.**
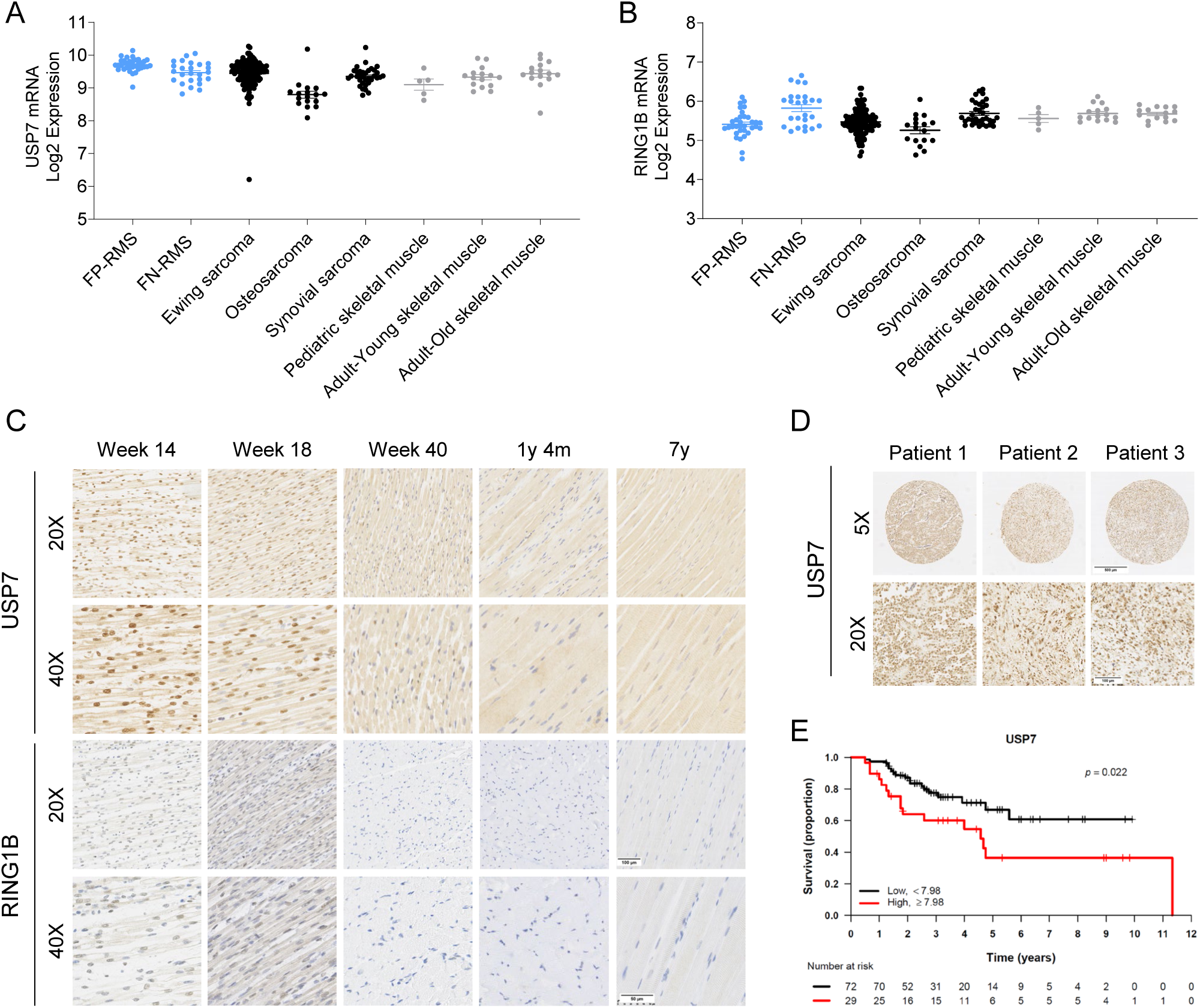
USP7 is overexpressed in FP-RMS and has prognostic value. **A.** USP7 mRNA expression from publicly available GEO databases (Affymetrix U133plus2.0 array) of pediatric tumors. Each dot represents a different sample. Sample sizes per group are as follows: RMS (n = 58), Ewing Sarcoma (n = 142), Osteosarcoma (n = 17), Synovial Sarcoma (n = 34). Source Data is available in Supplementary Table 1. **B.** RING1B mRNA expression from publicly available GEO databases (Affymetrix U133plus2.0 array) of pediatric tumors. **C.** Immunohistochemistry (IHC) of USP7 and RING1B in 5 out of 18 representative samples of muscle at different differentiation timepoints. From left to right: 14-weeks embryo, 18-weeks embryo, 40-weeks embryo, 1 year and 4 months child and a 7-year-old child. Images were acquired at 20× (top) and 40× (bottom) magnifications using a Leica CTR5000 microscope. **D.** IHC of USP7 in three TMA representative samples from Hospital Sant Joan de Déu. Images were acquired at 5X (top) and 20X (bottom) magnifications using a Leica CTR5000 microscope. **E.** Survival of RMS patients comparing high (≥ 7.98, n = 29, red) versus low (< 7.98, n = 72, black) levels of USP7 (p = 0.022).

To characterize the function of USP7 and RING1B in the RMS cell of origin context, we performed immunohistochemistry (IHC) staining in 18 human skeletal muscle samples at different stages of differentiation, ranging from a 14-week-old embryo to a 7-year-old child. We observed that nuclear staining of both USP7 and RING1B was progressively lost upon muscle differentiation with the greatest decrease shown in the 40-week embryo (**Figure 1C**). In the case of USP7, the remaining positive nuclei corresponded to satellite cells (muscle stem cells with flat nucleus located adjacent to muscle fibers), while a minimal staining of USP7 was retained in the cytoplasm of muscle fibers. All these data suggests that USP7 and RING1B are associated with the proliferative and undifferentiated state of precursor cells in the muscle differentiation process.

Next, we explored the expression of USP7 and RING1B in tumors using a tissue microarray (TMA) containing 26 RMS samples from patients treated at the Hospital Sant Joan de Déu. We observed uniform high levels of USP7 among all RMS samples, mainly localized in the nucleus but also present in the cytoplasm of tumor cells (**Figure 1D**), while RING1B showed variable levels (**Supplementary Figure 1C**). To explore whether USP7 might have prognostic significance in RMS, we reanalyzed the gene expression data of 101 RMS samples reported by Williamson et al. (E-TABM-1202)^6^. Upon data normalization using the Robust Multi-Array Average (RMA) method, we performed hierarchical clustering and samples grouped into FN-RMS or FP-RMS as expected (**Supplementary Figure 1D; Supplementary Table 2**). The survival analysis based on the presence or absence of the *PAX3::FOXO1* or *PAX7::FOXO1* fusion oncogenes showed that FP-RMS exhibit worse prognosis than FN-RMS in this patient cohort, concordant with the literature (**Supplementary Figure 1E**). These results established this cohort as representative of RMS, reproducing the main characteristics of FP-RMS and FN-RMS tumors. Survival curves of patients displaying high or low *USP7* or *RING1B* expression revealed that *USP7* (probe 201498_at) had prognostic value in RMS, with high expression of *USP7* associating with worse prognosis (p = 0.0022) (**Figure 1E**), while *RING1B* (probe 205215_at) had no prognostic significance (p = 0.13) (**Supplementary Figure 1F**). No association was observed between *USP7* levels and the presence of the fusion oncogene (p = 0.11), establishing that high *USP7* expression constitutes a negative prognostic factor for RMS patients independent of the fusion status. These findings highlight USP7 as a promising candidate for more in-depth studies to further elucidate its biological role and therapeutic relevance.

### USP7 and RING1B colocalize with PAX3::FOXO1 at active enhancers

Previous data indicate that USP7 modulates transcriptional activity and promotes gene expression in SySa and leukemia^25,28^. To gain a deeper understanding of the role of USP7 in FP-RMS, and to define its genome-wide occupancy together with the PRC1 catalytic subunit RING1B and the fusion oncoprotein PAX3::FOXO1, we performed chromatin immunoprecipitation followed by sequencing (ChIP-seq). We selected three PAX3::FOXO1 positive models for analysis: two cell lines (RH30 and RH4) and one primary model (HSJD-ARMS-006). Differential binding analysis (DiffBind)^41^ was performed on the peaks identified independently in two replicates for each protein to define a consensus set of common peaks **(Supplementary Figure 2A).** Additionally, to strengthen the robustness of downstream analyses, we focused on the TOP-5000 highest-scoring peaks of each protein. Similar trends are observed when working with full sets of peaks, although differences between groups on each comparison are less pronounced. Due to the lack of commercial antibodies against the fusion oncoprotein, we generated a PAX3::FOXO1 antibody (Innovagen AB, Sweden) using the peptide sequence of the published fusion region, thereby avoiding the detection of the wild-type form of PAX3 and FOXO1^8,42^. For antibody validation, we performed ChIP-seq in chromatin from the FN-RMS cell line RD, where we confirmed that no signal was detected, compared to positive signals in FP-RMS models (**Supplementary Figure 2B**).

Genomic distribution of peaks showed that PAX3::FOXO1 displayed a major preference for enhancers (introns and intergenic regions) than USP7 and RING1B, that showed greater enrichment at promoters despite also being distributed across genic and intergenic regions (**Figure 2A**). Classification of the genomic distribution of the peaks into active and poised enhancers and promoters, based on previously published histone marks in RH4 cells^8^, showed that PAX3::FOXO1, USP7 and RING1B were mainly enriched in transcriptionally active regions (**Supplementary Figure 2C**). MEME motif analysis of the TOP-5000 peaks for each protein revealed common motifs such as the paired domain (PD) binding motif of PAX3::FOXO1, the enhancer-box motif (E-box) and the Polycomb binding motif^43,44^, supporting the colocalization of these three proteins (**Figure 2B**). Consistent with the motif analysis, we confirmed that PAX3::FOXO1 binding sites, highly decorated with the active enhancer histone mark H3K27ac, were also enriched in USP7 and RING1B across all FP-RMS models (**Figure 2C, Supplementary Figure 2D**). We next intersected the TOP-5000 peaks of PAX3::FOXO1, USP7 and RING1B in RH30 and found that approximately 17% of peaks were shared among the three proteins, while 26% of PAX3::FOXO1 peaks overlapped with USP7 and nearly 50% with RING1B. Similar results were observed in RH4 and HSJD-ARMS-006, suggesting the presence of the PRC1.1 complex at these sites (**Figure 2D**).

**Figure 2.**
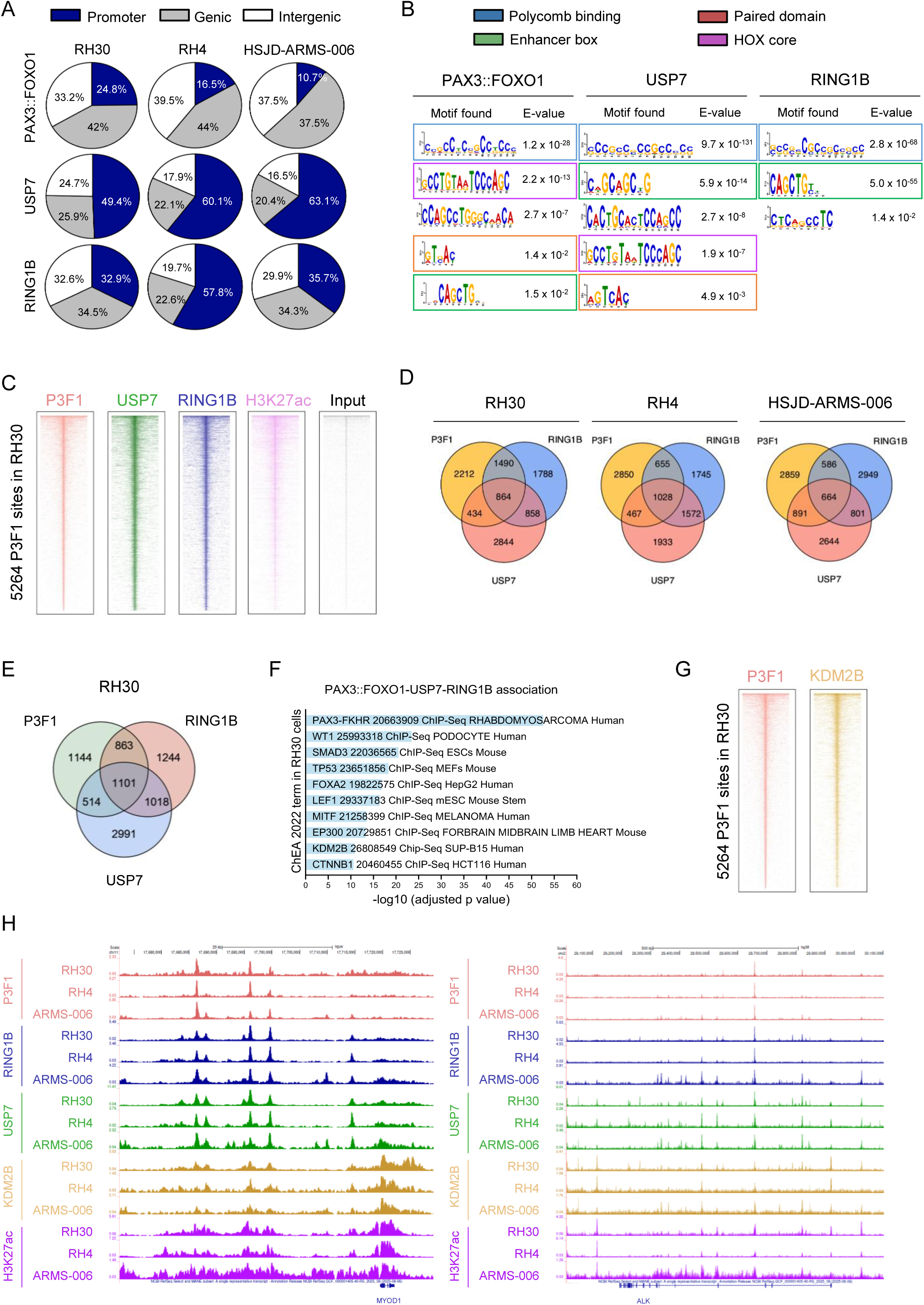
USP7 and RING1B colocalize with PAX3::FOXO1 at active enhancers. **A.** Pie chart showing genomic distribution of PAX3::FOXO1, USP7 and RING1B TOP-5000 peaks in RH30, RH4 and HSJD-ARMS-006. The three areas are promoters (blue), intragenic (grey) and intergenic (white) regions. **B.** Significant DNA binding motifs (E-value < 0.05) found by MEME analysis of PAX3::FOXO1, USP7 and RING1B in RH30 cells. Boxes highlight the following binding motifs: paired domain (PD) of PAX3::FOXO1 (red), Polycomb (blue), Hox core (purple), and enhancer box (E-box, green). DNA binding motifs are ordered by E-value. **C.** Heatmap of PAX3::FOXO1 (P3F1), USP7, RING1B, H3K27ac and negative control (input) ChIP-seq signals at PAX3::FOXO1 peaks, normalized by the total number of mapped reads in RH30. **D.** Venn-diagram depicting the overlap of PAX3::FOXO1, USP7 and RING1B peaks in RH30, RH4 and HSJD-ARMS-006 cells. **E.** Venn diagram depicting the overlap of genes associated with PAX3::FOXO1, USP7 and RING1B peaks in RH30 cells. **F.** Bar chart representing the top ten ChEA 2022 enriched categories of genes associated with PAX3::FOXO1, USP7 and RING1B common peaks in RH30 cells. Bar length indicates the -log10 (adjusted *p* value). All shown pathways have a *p* value < 0.05. Y axis shows the top 10 enriched pathways provided by EnrichR tool^45,46^. **G.** Heatmap of PAX3::FOXO1 and KDM2B ChIP-seq signals at PAX3::FOXO1 peaks, normalized by the total number of mapped reads. **H.** UCSC ChIP-seq signal tracks, normalized by the total number of mapped reads, for PAX3::FOXO1 (P3F1, red), RING1B (blue), USP7 (green), KDM2B (orange), and H3K27ac (pink) in RH30, RH4 and HSJD-ARMS-006 for MYOD1 (left) and ALK (right) genomic regions.

Due to the high colocalization between PRC1.1 subunits and PAX3::FOXO1, we identified genes located within 10 kb upstream or downstream of regions simultaneously occupied by these three proteins in RH30 cells, yielding 1101 genes (**Figure 2E**). Among these, we identified several well-known targets of the oncogene involved in FP-RMS tumorigenesis, including *ALK*, *FGFR4*, *FOXF1*, *JARID2*, *MYCN*, *MYOD1* and *MYOG*. Next, we performed a functional annotation of these genes with EnrichR (ChEA-2022 library)^45–47^ which revealed strong enrichment for genes identified in published ChIP-seq datasets for PAX3::FOXO1, p53 and p300, among others (**Figure 2F**; **Supplementary Figure 2E**). Interestingly, KDM2B, another defining subunit of the PRC1.1 complex, emerged as a significant match. To investigate this relationship further, we performed ChIP-seq for KDM2B in FP-RMS models. We observed significant colocalization of KDM2B with PAX3::FOXO1 peaks in RH30 cell line (**Figure 2G**), although KDM2B showed a preferential enrichment at promoters (**Supplementary Figure 2F**). Of the 1,101 genes associated with enhancers co-occupied by PAX3::FOXO1, USP7, RING1B and H3K27ac, 587 also overlapped with KDM2B peaks, including well known PAX3::FOXO1 targets such as *MYOD1* and *ALK* (**Figure 2H; Supplementary Figure 2G**). In contrast, the PRC1 catalytic subunit RING1A did not colocalize at PAX3::FOXO1 target sites, indicating that RING1B but not RING1A participates in the regulation of these active sites in FP-RMS (**Supplementary Figure 2H**). Collectively, these results support a fundamental role for PRC1.1 in shaping the epigenetic regulatory program of FP-RMS.

### USP7 downregulation reduces tumorigenic capacity of FP-RMS cells

To elucidate the contribution of USP7 to FP-RMS transcriptional programs and tumorigenesis, and following unsuccessful attempts to generate a CRISPR-Cas9 knock-out (KO) model, we depleted USP7 in RH30 and RH4 cell lines using a shRNA-mediated knock-down (KD) approach. The inability to generate a USP7 KO may reflect that the gene is essential for RMS survival, consistent with data from the Dependency Map (DepMap) portal. To evaluate the contribution of the PRC1 core, RING1B KO was successfully generated in RH30 cells. The decrease in USP7 and RING1B protein levels was validated by western blot (**Supplementary Figures 3A-B**). Characterization of the effects of USP7 and RING1B downregulation on other PRC1.1 components in RH30 cells revealed that KDM2B was decreased upon USP7 and sgRING1B #1 depletion, while a slight upregulation was observed in sgRING1B #2 (**Supplementary Figure 3B**). The reduction of KDM2B protein levels may be explained by the presence of USP7, RING1B and KDM2B peaks at the KDM2B promoter (**Supplementary Figure 3C**). Given the colocalization of USP7 and RING1B with PAX3::FOXO1, we aimed to determine whether these PRC1.1 subunits regulate the transcription of PAX3::FOXO1 target genes. RT-qPCR revealed that USP7 KD moderately reduced the expression of *RING1A* and *RING1B;* downregulated the core regulatory circuit (CRC) genes *MYOD1* and *MYCN* but not *MYOG;* and significantly downregulated PAX3::FOXO1 target genes *ALK, FGFR4, FOXF1, IGF1R* and *JARID2* (**Figure 3A**). Consistently, RING1B KO strongly downregulated CRC genes as well as PAX3::FOXO1 targets, besides a moderate decrease in *USP7* expression (**Supplementary Figure 3D**), supporting a PRC1.1 role in the oncogenic program of PAX3::FOXO1.

**Figure 3.**
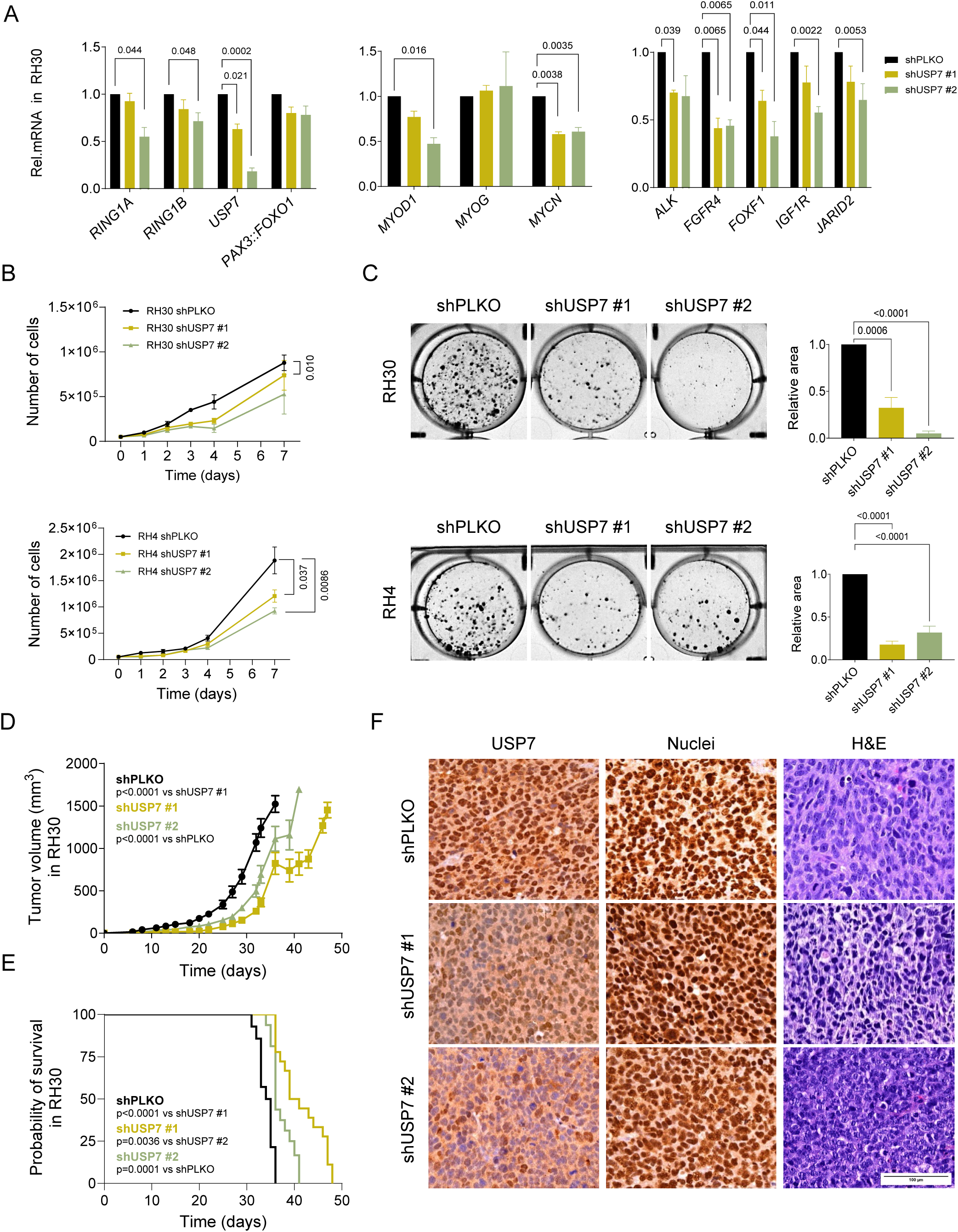
USP7 downregulation reduces tumorigenic capacity of FP-RMS cells. **A.** RT-qPCR determination of mRNA relative expression of PRC1.1 subunits genes (*RING1A, RING1B,* and *USP7), PAX3::FOXO1,* CRC genes *(MYOD1, MYOG, MYCN),* and oncogene targets *(ALK, FGFR4, FOXF1, IGF1R* and *JARID2)* in shUSP7 #1 and #2 compared to control in RH30 cells. TBP was used as a housekeeping gene for normalization. Significant p values are shown in the graph. **B.** Cell counts of RH30 and RH4 shPLKO, shUSP7 #1 and #2 for 7 days. P values are indicated when significant. **C.** Representative replicates of colony formation capacity in RH30 and RH4 USP7 KD cells. Quantification and p values of all replicates for RH30 USP7 KD colony formation capacity and RH4 are shown. **D.** Tumor volume curve in xenografts established by subcutaneous injection of RH30 shPLKO, RH30 shUSP7 #1 and shUSP7 #2. Tumors were eliminated when over-passed 1,500 mm^3^. **E.** Kaplan-Meier survival curves of RH30 shPLKO, RH30 shUSP7 #1 and RH30 shUSP7 #2. **F.** IHC of USP7 on sections of tumors excised from each group. Hematoxylin and Eosin (H&E) and human nucleus (nuclei) were used as a control of tumorigenic cells. Images were acquired at 40× magnifications using a Leica CTR5000 microscope.

Several markers of terminal muscle differentiation have been described as Myosin Heavy Chain 1 (*MYH1*) and Myogenic Factor 5 (*MYF5*). *MYH1* encodes a contractile motor protein characteristic of fast skeletal muscle fibers and is commonly used as a marker of muscle differentiation. *MYF5*, despite being an early transcription factor in the myogenic program, has been reported to be mutually exclusive with MYOD1 in RMS, suggesting reprogramming of the myogenic program or a compensatory mechanism^48,49^. Interestingly, these two markers were highly upregulated upon USP7 KD and RING1B KO (**Supplementary Figures 3E-F**). No peaks of RING1B, USP7, KDM2B and PAX3::FOXO1 were found in the promoters or in proximal genomic regions near the transcription start sites (TSSs) of *MYH1* and *MYF5*, suggesting indirect upregulation. Altogether, these findings suggest that both USP7 and RING1B might be contributing to the maintenance of the oncogenic transcriptional programs in FP-RMS by regulating PAX3::FOXO1 target genes, while their depletion promotes myogenic differentiation. Next, we investigated the potential role of USP7 and RING1B in FP-RMS tumorigenesis. Growth curves demonstrated that USP7 and RING1B downregulation significantly decreased cell proliferation under most conditions in RH30 and RH4 cell lines (**Figure 3B**; **Supplementary Figure 3G**). In line with these results, we observed that shUSP7 #1 and #2 significantly reduces the colony formation capacity in RH30 (p = 0.0006 and < 0.0001, respectively) and RH4 (p < 0.0001, both) compared to controls (**Figure 3C**). Consistently, RING1B KO significantly decreased the colony formation capacity of RH30 cells (**Supplementary Figure 3H;** sgRING1B #1; p =0.0007 and #2; p = 0.0013). To evaluate whether USP7 has a role in tumor growth *in vivo*, we injected RH30 shPLKO control cells and RH30 shUSP7 #1 and #2 subcutaneously into mice. We observed that shUSP7 #1 and #2 exhibited a significantly tumor growth delay compared to shPLKO control group (p < 0.0001 for both comparisons) (**Figure 3D**). Moreover, both USP7 KD (#1 and #2) cells showed an improved survival extended from day 36 to day 41 and 48, respectively, when compared to shPLKO condition (shUSP7 #1; p < 0.0001 and #2; p = 0.0001) (**Figure 3E**). IHC analysis of tumors confirmed reduced levels of USP7 as causative of the decreased growth (**Figure 3F**). Altogether, these data suggest that both USP7 and RING1B contribute to FP-RMS tumorigenesis.

### USP7 inhibitor P22077 reduces FP-RMS tumor growth *in vivo*

To investigate whether PRC1.1 inhibition through USP7 targeting might become a novel therapeutic strategy for FP-RMS, we selected the widely used USP7 inhibitor P22077^37^. To that end, we performed viability assays in a panel of different FP-RMS cell lines and primary cultures including RH30, RH4, RH28, and HSJD-ARMS-006. Non-RMS cell lines HeLa and LHCN-M2 were included as controls. We observed that all FP-RMS cell lines responded to P22077 in a dose-dependent manner (**Figure 4A**), with RH28 and HSJD-ARMS-006 showing the highest sensitivity, whereas HeLa cells did not respond to P22077 (**Figure 4B**). The myoblast cell line LHCN-M2 exhibited a pronounced cytotoxic response at 12 μM, a concentration above the IC50 range observed in FP-RMS. Since USP7 has been described to deubiquitinate H2BK120ub^50^, we assessed the levels of this histone modification as a marker of USP7 inhibition. An increase in H2BK120ub confirmed the inhibition of USP7 enzymatic activity in RH30 and RH28 cells lines (**Supplementary Figure 4A**). To evaluate the potential cytotoxic effect of USP7 inhibition with P22077, we assessed cPARP and observed increased levels across all cell lines and FP-RMS ARMS006 when treated at their IC50 (**Figure 4C**). Indeed, cPARP increased in a dose-dependent manner following P22077 treatment in the RH30 cell line (**Supplementary Figure 4B**). To confirm the specificity of USP7 inhibition, we selected the commercial USP7 PROTAC U7D-1 that specifically targets and degrades USP7 via the proteasome^51^. We validated that U7D-1 downregulates USP7 levels in RH30 cells at 0.5 μM, concomitant with an increase in cPARP (**Supplementary Figure 4C**). Considering the role of USP7 in protein stabilization, we assessed the protein levels of PRC1.1 subunits. We observed a slight decrease in KDM2B protein levels, in agreement with our observations in USP7 KD, while no other PRC1.1 subunits were altered (**Supplementary Figure 4D**). Unaltered proteins levels of USP7 are consistent with previous studies showing that P22077 impairs USP7 activity by binding to its enzymatic domain without inducing degradation. Although USP7 inhibition does not alter the global protein levels of the histone marks H3K27me3, H3K27ac or H2AK119ub, we detected an increase in γH2AX levels in all three FP-RMS cell lines upon P22077 treatment (**Supplementary Figure 4E**). Moreover, while PAX3::FOXO1 levels were maintained, USP7 inhibition increased p21 levels, consistent with previous publications suggesting cell cycle arrest and the presence of DNA damage, as indicated by increased γH2AX levels (**Supplementary Figure 4F**)^52^.

**Figure 4.**
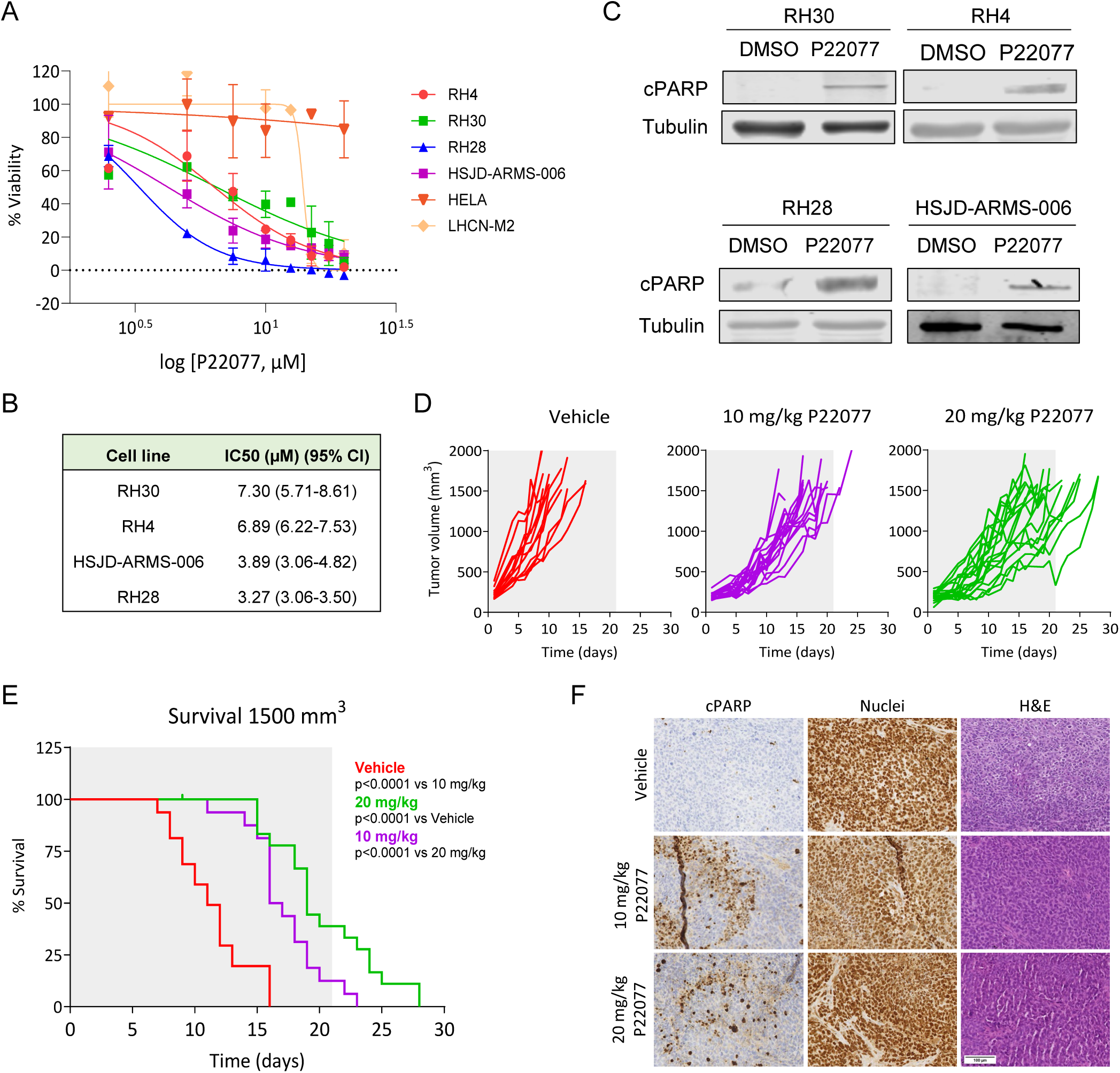
USP7 inhibitor P22077 reduces FP-RMS tumor growth *in vivo*. **A.** Cell viability assays showing the trend line of concentration-dependent cytotoxicity of P22077 treatment in FP-RMS cell lines RH30, RH4, RH28 and HSJD-ARMS-006 and control cell lines HELA and LHCN-M2 at 72 h. Percentage of cell viability is normalized to the average viability of the drug vehicle (DMSO). X-axis represents the Log10 of P22077 expressed in μM. Plotted is the mean normalized viability of three replicates ± the SEM. **B.** IC50 values of P22077 for FP-RMS cell lines calculated using Graphpad Prism. **C.** Western blot showing cPARP levels of RMS cells treated with P22077 at their IC50 concentration. Tubulin was used as a loading control. **D.** Tumor volume curves in xenografts injected with RH30 cells and treated with vehicle (red), 10 (purple) and 20 (green) mg/kg of P22077. Each line represents an individual tumor. Grey shadow represents the duration of the treatment. **E.** Survival curves of RH30 vehicle (red), 10 mg/kg of P22077 (purple) and 20 mg/kg of P22077 (green). **F.** IHC staining of cPARP in vehicle, 10 mg/kg and 20 mg/kg P22077 groups. Hematoxylin and Eosin (H&E) and human nucleus (nuclei) were used as a control of tumorigenic cells. Samples were collected after 5 doses and 2.5 h after the last dose of P22077.

To further demonstrate the effect of USP7 inhibition in FP-RMS, we injected subcutaneously RH30 cells into mice. After random distribution into three groups when tumors reached 150–300 mm^3^, mice were treated intraperitoneally six days per week for three weeks with two doses of P22077 (10 or 20 mg/kg) or vehicle. We observed that tumors in the P22077-treated groups grew significantly slower than the vehicle group, with the 20 mg/kg group showing the strongest delay (p < 0.0001, **Figure 4D**). However, differences in growth rates between both treated groups were not significant (p = 0.1310). Survival was significantly improved upon P22077 treatment from day 16 in the vehicle group to days 23 and 28 in 10 mg/kg and 20 mg/kg of P22077 groups, respectively (p < 0.0001, both) (**Figure 4E**). In this case, differences in survival were significant between both treated groups (p < 0.0001). Notably, we did not observe toxicity in any groups as assessed by body weight loss and animal welfare deterioration (**Supplementary Figure 4G**). To characterize the effects of P22077 on tumors *in vivo*, we euthanized two random animals of each group after five doses of P22077. We observed an increase in cPARP by IHC in both treated groups with evidence of apoptosis in the tumor (**Figure 4F**). These results support P22077 as a potential therapeutic strategy for FP-RMS.

### USP7 inhibition reduces the expression of PAX3::FOXO1 targets in FP-RMS

USP7 inhibition reduces FP-RMS cell viability both *in vitro* and *in vivo*. To elucidate the impact of USP7 inhibition on gene expression following P22077 treatment, we performed RNA sequencing (RNA-seq, three biological replicates) using three FP-RMS cell lines (RH4, RH30 and RH28) comparing the control condition (DMSO) and P22077-treated conditions at the IC50 for each cell line. Principal Component Analysis (PCA) revealed clear segregation between controls and the USP7 inhibitor-treated triplicates within each cell line (**Supplementary Figure 5A**). Notably, the RH30 and RH28 cell lines clustered closely together in the PCA analysis. Next, we performed differential gene expression analysis using DESeq2^53^ (adjusted P value < 0.1) to compare treated and control conditions. Although RH30 and RH28 showed higher percentages of downregulated genes, supporting USP7 as a transcriptional activator in FP-RMS, upregulated genes exhibited greater fold-change magnitudes (**Supplementary Figure 5B; Figure 5A**). Functional analysis of the biological pathways in the differentially upregulated genes reported significant enrichments on pathways related to muscle processes such as muscle organ development, muscle tissue development, muscle cell development and differentiation (**Figure 5B; Supplementary Figure 5C**). These results indicate that USP7 inhibition induces muscle differentiation in FP-RMS cell lines. On the other hand, downregulated genes (log2FC< 0.5 and FDR < 0.05) according to the ChEA 2022 library (EnrichR) revealed a significant enrichment of PAX3::FOXO1 and MYC targets in RH30 and RH28 cells (**Figure 5C; Supplementary Figure 5D**). To gain insight on this aspect, we performed GSEA enrichment analysis of the RH30 full transcriptome of both conditions ranked by the DESeq2 P22077/DMSO stat value using as gene set the TOP5000 PAX3::FOXO1 targets identified in our previous ChIP-seq. A negative enrichment was reported in the GSEA plot (NES −1.58, FDR q-value 0.0, **Figure 5D**) indicating a dual transcriptional regulatory role of USP7 and PAX3::FOXO1. Indeed, the same GSEA comparison for the PAX3::FOXO1, RING1B, USP7 and KDM2B common targets identified by ChIP-seq experiments as gene set reproduced a similar negative enrichment (NES −1.62, FDR q-value 0.0, **Figure 5E**). In these gene signatures we found interesting known regulators as *ALK, FGF8, FGFR2, FGFR4, FOXF1, IGF2, JARID2, MEOX1, MYCN* and *SOX5*.

**Figure 5.**
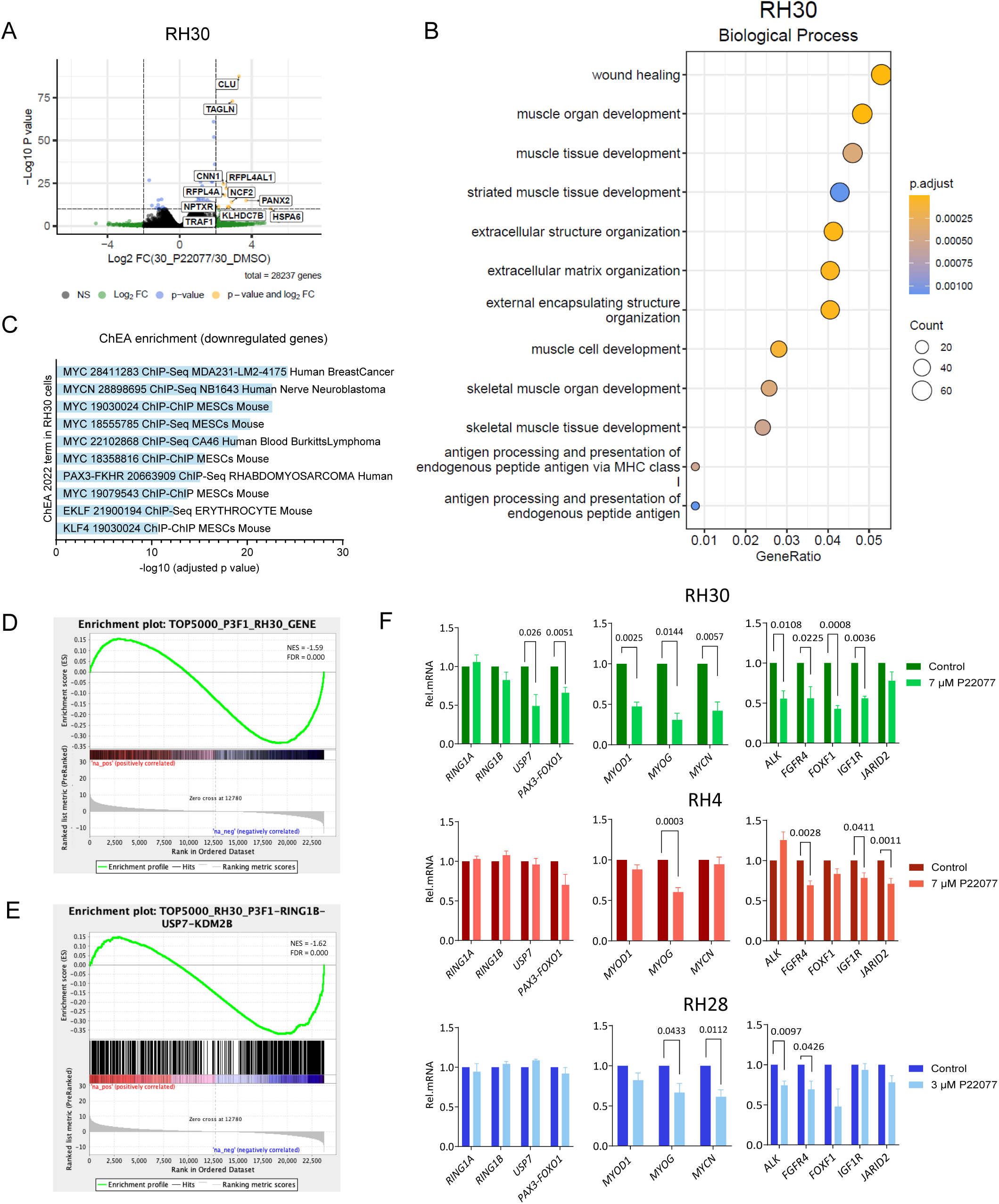
USP7 inhibition reduces the expression of PAX3::FOXO1 targets in FP-RMS. **A.** Volcano plot showing RNA-seq differential gene expression between DMSO and P22077 conditions. **B.** Bubble chart depicting the top 12 enriched gene ontology (GO) Biological Processes of upregulated genes upon USP7 inhibition in RH30 cells. Categories are ranked by the fraction of the gene set belonging to this function. Circle size represents the number of genes on the list belonging to this function. Circle color denotes the adjusted p value of each enrichment. **C.** Bar chart representing the top ten ChEA 2022 enriched categories of downregulated genes by RNA-seq in RH30 upon USP7 inhibition with P22077. **D.** GSEA enrichment plot showing the distribution of genes that are targets of PAX3::FOXO1 across the list of genes ranked by the stat value obtained from DESeq2 analysis between DMSO and P20077 conditions. **E.** GSEA enrichment plot showing the distribution of genes that are targets of PAX3::FOXO1, USP7, RING1B and KDM2B across the list of genes ranked by the stat value obtained from DESeq2 analysis between DMSO and P20077 conditions. **F.** RT-qPCR determination of mRNA relative expression of PRC1.1 subunits genes, *PAX3::FOXO1,* CRC genes and oncogene targets in DMSO versus P22077 conditions in RH30 (green), RH4 (red) and RH28 (blue). TBP was used as a housekeeping gene for normalization. Significant p values are shown in the graph.

To further validate the potential role of USP7 in regulating oncogenic gene expression in FP-RMS, we explored differences in the expression of CRC genes and selected key targets of PAX3::FOXO1 previously analyzed in the USP7 KD and RING1B KO experiments. While *RING1A* and *RING1B* did not change upon P22077 treatment, *USP7* and *PAX3::FOXO1* expression was downregulated in RH30 cells (**Figure 5F**). Consistent with the RNA-seq data, three genes of the FP-RMS core regulatory circuit (*MYOD1*, *MYOG* and *MYCN*) and PAX3::FOXO1 target oncogenes were significantly downregulated upon USP7 inhibition, reinforcing the role of USP7 sustaining the oncogenic program of PAX3::FOXO1 through binding to their SEs. Comparable results were obtained in two other PAX3::FOXO1 positive cell lines (RH28 and RH4) as well as in the primary FP-RMS cell culture (HSJD-ARMS-006) (**Figure 5F; Supplementary Figure 5E**). USP7 degrader U7D-1 further confirmed the downregulation of a subset of these targets (**Supplementary Figure 5F**). Overall, these data support an essential role of USP7 in the chromatin remodeling process infringed by PAX3::FOXO1 and MYCN in FP-RMS.

### USP7 inhibition induces myogenic differentiation

To further explore the effect of USP7 inhibition on myogenesis-related pathways (**Figure 5B**), we performed Gene Set Enrichment Analysis (GSEA) using the Molecular Signatures Database (gene set Hallmark_Myogenesis) on the differentially expressed genes (DEGs) identified in our previous RNA-seq comparison between P22077 and DMSO conditions. Heatmap analysis demonstrated increased expression of genes related to myogenesis in RH30 (**Figure 6A**), with genes such as *MYOM1* and *CLU*, significantly upregulated (**Figure 6B**). Our analysis confirmed the effect of USP7 inhibition to induce myogenesis, as evidenced by positive enrichment of genes within this signature, suggesting activation of muscle terminal differentiation in RH30, RH4 and RH28 **(Figure 6C; Supplementary Figure 6A**). MYCN contributes to the maintenance of an undifferentiated state in RMS, whereas its downregulation has been associated with the induction of myogenic differentiation in *in vitro* models^54^. Strikingly, when we performed a GSEA analysis interrogating the P22077/DMSO transcriptomic comparison with the MYC hallmark signature, we identified a more pronounced negative enrichment upon P22077 treatment (**Figure 6D**), consistent with a progression toward a myogenic state. We next selected the muscle differentiation markers MYH1 and MYF5 to evaluate the induction of myogenesis upon P22077 treatment, observing an increase at both RNA expression and protein levels in the different cell models (**Figure 6E-F; Supplementary Figures 6B**). Furthermore, we validated the differential expression of selected myogenic genes identified by RNA-seq using RT-qPCR (**Supplementary Figure 6C**). In agreement, the USP7 specific degrader U7D-1 increased the expression of differentiation markers *MYH1* and *MYF5* (**Supplementary Figure 6D)**. Altogether, these results support that the PRC1.1 complex regulates PAX3::FOXO1 target genes and the myogenic state of FP-RMS cells, representing its inhibition a potential therapeutic strategy inducing FP-RMS differentiation.

**Figure 6.**
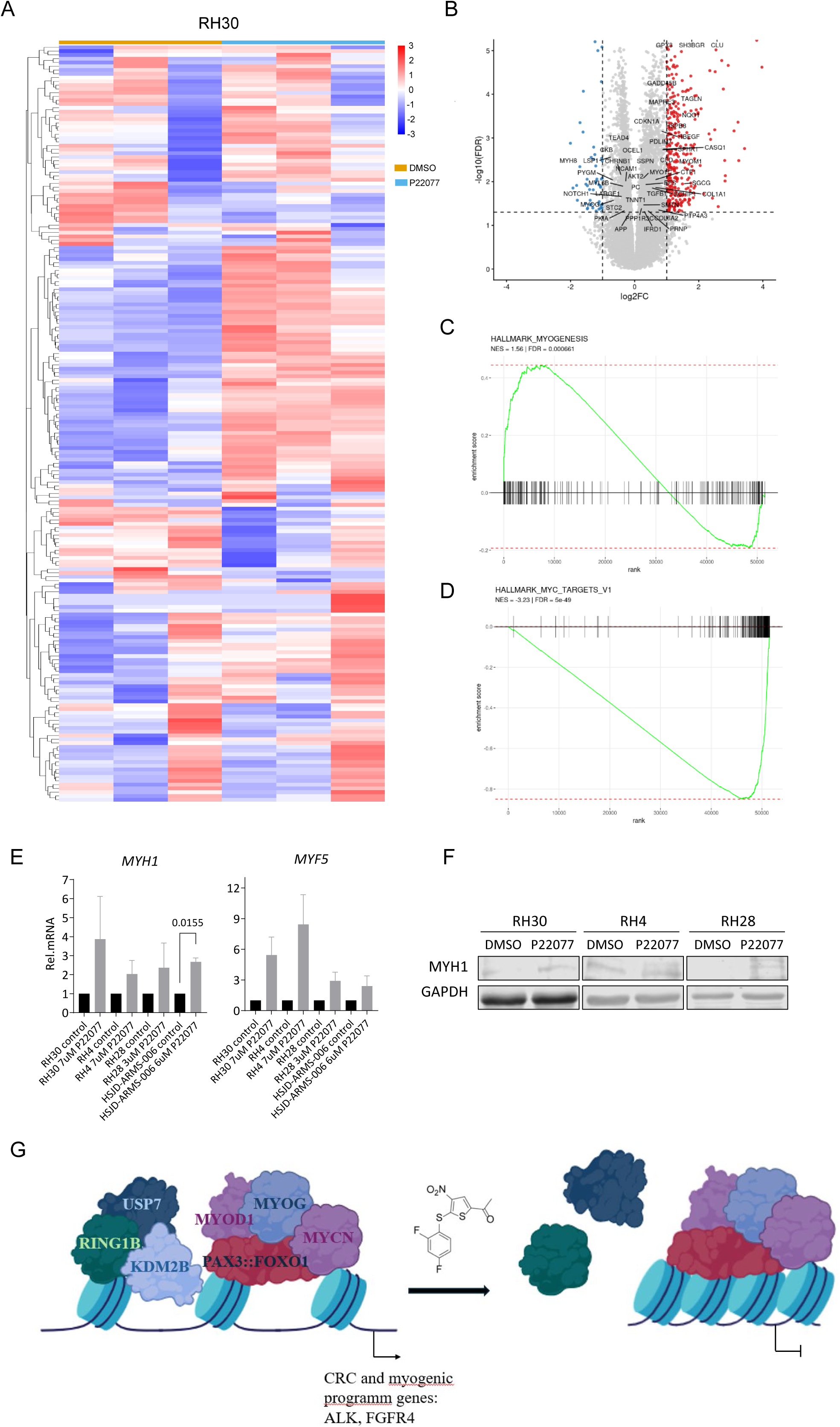
USP7 inhibition induces myogenic differentiation in FP-RMS. **A.** Heatmap showing the expression pattern of genes from the Molecular Signatures Databases (MSigDB, gene set Hallmarks_Myogenesis) in RH30. Gene expression values are displayed as row-scaled normalized expression (z-scores), where red indicates higher expression and blue indicates lower expression relative to the mean across samples. Columns represent individual biological replicates and rows represent genes. **B.** Volcano plot showing RNA-seq-derived genes from the Hallmark_Myogenesis (MSigDB) between DMSO and P22077 conditions **C.** GSEA **e**nrichment plot for the Hallmark_Myogenesis (MSigDB). The green curve represents the running enrichment score (ES). Black vertical bars indicate the positions of individual genes from the gene set within the ranked dataset. **D.** GSEA **e**nrichment plot for the Hallmark_MYC_targets (MSigDB). **E.** RT-qPCR determination of mRNA relative expression of *MYH1* (left) and *MYF5* (right) in RH30, RH4, RH28 and HSJD-ARMS-006 in control versus P22077 conditions. Cells were treated at their IC-50. TBP was used as a housekeeping gene for normalization. **F.** Western blot showing MYH1 levels upon P22077 treatment in RH30, RH4 and RH28 cells treated at their IC50 of P22077. GAPDH was used as a loading control. **G.** Schematic model in which USP7 inhibitor induces differentiation through disruption of PAX3::FOXO1-dependent enhancer reprogramming of genes sustaining undifferentiated state.

## DISCUSSION

PAX3::FOXO1 reprograms the epigenetic landscape of FP-RMS cells, generating specific transcriptional programs driven by oncogenic enhancers that represent a potential therapeutic vulnerability in these tumors^8,55,56^. Chromatin modifiers contributing to enhancer activation have emerged as key drivers of transcriptional reprogramming in fusion-driven pediatric sarcomas. In this scenario, we investigated the genome-wide contribution of the PRC1.1 subunit USP7, along with RING1B and KDM2B.

Survival analysis of retrospective cases identifies high USP7 expression levels as a candidate biomarker of poor prognosis. Although analyses in larger databases as well as prospective validation will be necessary to establish USP7 as a prognostic factor, the association with worse prognosis supports the importance of USP7 in RMS. Likewise, in neuroblastoma, USP7 is more abundant in patients with worse prognosis correlating with higher MYCN activity^35^. Moreover, SNAI2 regulation through USP7 participates in FN-RMS vasculogenic, proliferative, and migratory processes^57^. In agreement with DepMap and despite multiple attempts, we were unable to generate a stable USP7 KO in FP-RMS cell lines, highlighting the functional importance of USP7 and pointing to this PRC1.1 subunit as a vulnerability in RMS.

We identified USP7 and RING1B at PAX3::FOXO1 intergenic and intronic active enhancers decorated with H3K27ac. The differential role of RING1B at oncogenic enhancers has been previously observed in melanoma, breast cancer, leukemia, and EwS^21–24^. Here we confirm its presence at the transcriptionally active regions in FP-RMS, concomitant with the presence of USP7, which we now describe as playing a role at oncogenic translocation-driven enhancers for the first time. Given the specificity of USP7 for the PRC1.1 complex, our results suggest that this non-canonical PRC1 complex underlines the RING1B activity at these oncogenic enhancers. Nevertheless, the observed enrichment of USP7 and RING1B at active promoters compared to PAX3::FOXO1, suggests a broader role in transcriptional regulation. In contrast to recent publications that describe RING1A as an activator^58^, we do not observe RING1A enrichment at PAX3::FOXO1-driven active enhancers. These results suggest that RING1A may retain a specific role at repressed regions similar to what we previously described in EwS^24^.

We found that well-described targets of PAX3::FOXO1 are downregulated upon RING1B KO, USP7 KD or pharmacological inhibition of USP7. Some of these targets, such as *ALK*, *FGFR4, FOXF1*, *IGF1R* or *JARID2*, as well as *MYOD1*, *MYOG* and *MYCN* transcription factors from the CRC, are key players in FP-RMS tumorigenesis. Modulation of their expression by USP7 knockdown and inhibition might contribute to the decreased tumor volume and increased survival observed, supporting an oncogenic role for PRC1.1 in this context (**Figure 6G**). Thus, we show that an aberrant transcription factor such as PAX3::FOXO1 relies on USP7 to activate enhancers, promoting an altered gene expression profile that favors cell transformation. The reduction of colony formation capacity upon RING1B depletion supports PRC1.1 role in FP-RMS, in agreement with the oncogenic properties previously demonstrated by our group in EwS^24^. Our data confirm that PRC1.1 plays a key role in regulating tumorigenesis along with a fusion oncogene as previously described in SySa^25–27^.

Besides the role of USP7 in regulating the p53 axis, USP7 has been described to promote tumorigenesis by deubiquitinating and stabilizing multiple oncogenic factors. In leukemia, PRC1.1 chromatin binding depends on USP7 deubiquitinase activity and its inhibition induces loss of PRC1.1 occupancy through the disruption of the complex^28^. In agreement, USP7 has been described to interact with and stabilize KDM2B^29,59^. These data are supported by a decrease in KDM2B levels upon USP7 inhibition and knockdown (**Figure 6G**). In neuroblastoma, USP7 expression induces deubiquitination and subsequent stabilization of MYCN, supporting USP7 inhibition as a potential therapy for *MYCN* amplified tumors^35^. In the case of FP-RMS, MYCN has been defined as a component of the core regulatory circuit that maintains elevated expression of MYOD1, MYOG and PAX3::FOXO1^8^. Thus, the negative enrichment of MYCN targets upon USP7 inhibition indicates that USP7 promotes MYCN activity in FP-RMS, and its inhibition disrupts the maintenance of the oncogenic phenotype.

PRC1 participates in the myogenesis program repressing genes involved in differentiation. Transcriptome datasets of more than 400 human skeletal samples suggest that chemical inhibition of the PRC1 complex may induce muscle cell differentiation^38^. Nevertheless, myoblasts lacking RING1B failed to differentiate efficiently, supporting that loss of PRC1 prevents progression of myogenesis^13^. Whereas in muscle lineage cells PRC1 silences genes required for myogenic differentiation, in FP-RMS we describe PRC1.1 promoting transcription at active enhancers and promoters of PAX3::FOXO1 and MYCN, contributing to the elevated MYOD1 and MYOG expression. This differential behavior of PRC1 in the normal versus tumoral contexts may explain why inhibition of RING1B stimulates muscle differentiation in both scenarios. Accordingly, different subsets of PRC1 complexes can provide their activity in each context. RNA-seq experiments revealed an increase in myogenic differentiation markers and pathways in FP-RMS cells, alongside a strong decrease in PAX3::FOXO1 target gene expression upon PRC1.1 inhibition with P22077. The contribution of USP7 to the block in myogenic differentiation in FP-RMS likely involve multiple mechanisms. First, we hypothesize that the high levels of USP7 observed in fetal samples are consistent with elevated USP7 levels described in mouse pluripotent cells, where USP7 is required to maintain an undifferentiated state by repressing mesoendodermal lineage genes together with SRY-Box Transcription Factor 2 (*SOX2)*^60^. In addition, MYCN maintains transcriptional programs necessary for sustaining the undifferentiated myogenic state^61^. Similar to neuroblastoma, USP7-mediated stabilization of MYCN in FP-RMS may represent a conserved hijacked oncogenic mechanism. Thus, loss of MYCN activity upon USP7 inhibition may cooperate in promoting myogenic differentiation. Furthermore, the induction of differentiation markers with the PROTAC degrader U7D-1 supports the specificity of these observations.

Inhibition of USP7 has emerged as a promising preclinical strategy in multiple tumor types, including leukemia, neuroblastoma and other adult cancers, through regulation of PRC1.1 and/or the p53 pathway^28,36,37^. Ongoing preclinical studies are evaluating safety and efficacy, highlighting the potential to overcome chemoresistance and prevent tumor progression in multiple malignancies. In FP-RMS, our results support a model in which the PRC1.1 subunits USP7 and RING1B play pivotal roles in facilitating PAX3::FOXO1 transcriptional activation of target genes that shape the FP-RMS epigenetic landscape. These results identify novel therapeutic opportunities for RMS based on its epigenetic dependencies.

## MATERIAL AND METHODS

### Culture conditions and PAX3::FOXO1 antibody generation

FP-RMS commercial cell lines *PAX3::FOXO1* positive (RH30, RH4, RH28), *PAX7::FOXO1* positive (CW9019) and FN-RMS (RD) were cultured in RPMI 1640 media (Gibco®), supplemented with 10% FBS (Fetal Bovine Serum, Gibco®), 100 units/mL L-Glutamine and 100 μg/mL penicillin/streptomycin. Cell lines were grown in a humidified incubator at 37ᵒC under 5% CO_2_. When required, cells were split and diluted with Trypsin-EDTA solution (Gibco®). For long-term storage, cells were cryopreserved in liquid nitrogen in FBS supplemented with 10% DMSO. All RMS cell lines were kindly provided by the laboratory of Dr. Óscar Martínez-Tirado. HELA and HUVEC cell lines were acquired from the American Type Culture Collection (ATCC). LHCN-M2 cell line, kindly provided by Dr. Cecilia Jimenez-Mallebrera, was maintained in a medium consisting of DMEM high glucose (11995065, Gibco) and M-199 (31150022, Thermo Fisher Scientific) in a 3:1 ratio, supplemented with Glutamine 1%, Penicillin-Streptomycin (1%), FBS (15%), HEPES 0.02 M, Zinc Sulfate 0.03 μg/ml, B12 Vitamin 1.4 μg/ml, Dexamethasone 0.055 μg/ml, HGF 2.5 ng/ml and FGF 10 ng/ml.

Primary culture HSJD-ARMS-006 was generated from a patient-derived xenograft (PDX). A fresh biopsy obtained from a patient at Hospital Sant Joan de Déu was implanted and expanded in mice following informed consent and approval by the institutional ethics committee. When the implanted tumor achieved a size around 1,000 mm^3^, tumors were collected, mechanically dissociated, filtered with 0.45 μM CorningTM Sterile Cell Strainers (Fisher Scientific) and cultured in Tumorspheres Stem Media (TSM)^62^. All cell lines were authenticated by short tandem repeat profiling and routinely tested for Mycoplasma contamination, with all results remaining negative throughout the study.

To generate a PAX3::FOXO1 antibody for ChIP experiments, we obtained the peptide sequence of the fusion region from Cao L. et al., 2010^42^. We designed a polyclonal antibody with Innovagen, similarly to what was previously performed^8,42^.

### Patient samples

Biopsies from 18 human skeletal muscle samples and a tissue microarray containing 26 primary RMS tumors at diagnosis from the HSJD Biobank, integrated in the Spanish Biobank Network of ISCIII and in the Xarxa de Tumors de Catalunya, were used for experimental purposes in agreement with the ethical committee procedures. Following protocols received additional approval from Institutional Review Boards at Fundació Sant Joan de Déu and Hospital Sant Joan de Déu. Written informed consent was obtained from patients or their legal guardians before collection of samples.

### Plasmids and lentiviral infection

We generated an *USP7* knock-down in FP-RMS cell lines RH30 and RH4 following standard protocol^24^. Sequences targeting USP7 are described in Supplementary Table S6. The pLKO.1-puro vector (Sigma-Aldrich), harbouring a puromycin resistance gene was used. Lentiviruses were produced using HEK-293T packaging cells co-transfected with the MISSION pLKO.1-shRNA construct and three packaging plasmids (RSV, VSVG and RRE) using X-tremeGENE transfection reagent (Roche) diluted in OptiMEM (Gibco). Following 48 h incubation the supernatant was collected and added to RMS cells in the presence of 1 µg/mL polybrene (Sigma-Aldrich) to enhance transduction efficiency. After removal of the viral supernatant, cells were selected with 2 µg/mL puromycin. Selection was maintained until all non-transduced control cells were eliminated.

### CRISPR-Cas9 knock-out system

Gene knockout was performed using CRISPR-Cas9 ribonucleoprotein (RNP) complexes delivered by lipofection. The Gene Knockout Kit v2 (Bionova/Synthego) was used according to the manufacturer’s instructions. Cas9, Lipofectamin Cas9 Plus Reagent, Lipofectamine CRISPRMAX Transfection Reagent and sgRNA against *RING1B* or *USP7* were added to 0.5·10^5^ cells per condition resuspended in OptiMEM I Reduced Serum Medium. Sequences are described in Supplementary Table S6. Cells were seeded into 6-well plates, and CRISPR-edited cell pools were isolated 48 h later by limiting dilution in 96-well plates. Following 2–4 weeks of expansion, clones were sequenced using a 3500 Genetic Analyzer Hitachi (Applied Biosystems), and protein expression levels were assessed by western blot. The Inference of CRISPR Edits (ICE) analysis software was used to analyze the sequences of edited cells (clones with a model fit R^2^ > 0.6 were only considered). Clones carrying a mutated DNA sequence and lacking protein expression were considered valid KO clones and used for subsequent studies.

### Cell extract preparation and western blotting

Whole cellular extracts were prepared with RIPA buffer (150mM NaCl, 50mM Tris pH 8, 0.5% deoxycholic acid, 0.1% SDS, 1X NP-40) supplemented with EDTA-free protease inhibitor cocktail (Roche). Cell lysates were incubated 30 minutes at 4 °C and then centrifuged at 14.000 rpm for 15 minutes at 4 °C. Histone extracts of cultured cells were isolated using the EpiQuick Histone Extraction kit (Epigentek) following the manufacturer’s instructions. Protein quantification was done using Bradford reagent (Bio-Rad Protein Assay Dye Reagent, Bio Rad). 30–50 μg of whole protein extracts or 5 μg of histones were resolved by polyacrylamide gel electrophoresis and western blotting was performed using standard protocols. Incubation with primary antibodies was performed at 4°C overnight and LI-COR secondary antibodies that are detectable by near-infrared fluorescence were used for detection. The primary antibodies and dilutions are listed (see Supplementary Table S6). Secondary antibodies goat anti-rabbit and goat anti-mouse IRDye were diluted 1:10,000 and incubated for 1 h at room temperature to blotted membranes. Nitrocellulose membranes (Sigma-Aldrich) were scanned and visualized with an Odyssey CLx Infrared Imaging System at medium intensities.

### Cell viability and colony formation assays

IC50 calculations were determined in cell lines and primary cultures by cell viability assay. Cell lines were seeded in a concentration of 3.000 cells/well and primary cultures in a concentration of 30.000 cells/well in 96-well plates. Different doses of P22077 were added in serial dilutions and vehicle (DMSO) was included as a control. 10% of MTS reagent (G5421 - Promega®) per well was added 72 hours post treatment while protecting from light. Absorbance at 490 nm was measured using a Tecan plate reader after 2-4 hours of incubation. For viability calculation, GraphPad Prism 10 software was used using a non-linear regression analysis using a log (inhibitor) versus response and constrained to 100-0 viability to determine IC50 values.

Clonogenic assays were performed by seeding 0,0750·10^4^ cells/mL in 6-well plates and changing media every 2-3 days until visible colonies were grown. At endpoint, colonies were fixed with formalin 4% (PFA) and stained with crystal violet (2% crystal violet and 20% methanol in PBS 1X). Colony quantification was performed by area measurement using ImageJ/FIJI software.

### RNA extraction and RT-qPCR

Total RNA was isolated and purified using the RNeasy Mini Kit (Qiagen) according to the manufacturer’s protocol. Quantification of RNA samples was performed using Nanodrop 1000 spectrophotometer. Reverse transcription (RT) was performed with 1-μg of RNA sample converted to cDNA in a reaction catalyzed by a retrotranscriptase enzyme (M-MLV Reverse Transcriptase Promega). Random primers and RNase inhibitor (RNasin Plus RNase Inhibitor, Promega) were also added to the reaction. cDNA obtained was analyzed by qPCR using SYBR Green PCR Master Mix (ABI). cDNA was amplified with specific oligonucleotides (see Supplementary Table S6). Each cDNA sample was run in triplicate, and gene expression levels were analyzed using the 7500 Fast PCR instrument (Applied Biosystems). To compare between different conditions, relative quantification of each target was normalized to a housekeeping gene. Data were analyzed using the comparative 2^-ΔΔct^ method.

### Microarray gene expression

To analyze the expression of RING1B and USP7 in RMS and other sarcomas and healthy tissues we selected public GO datasets for pediatric tumor transcriptomes generated with the platform U133 plus2.0, Affymetrix: GSE59880, GSE27661, GSE139400, GSE8840, GSE69698, GSE67326, GSE14827, GSE17679, GSE20196, GSE37371 and GSE66533. This database includes 58 RMS patient samples (25 FN-RMS and 33 FP-RMS) and 11 RMS cell lines (4 FN-RMS and 7 FP-RMS) among a collection of samples from other pediatric sarcomas (142 EwS, 17 OS, 34 SynS). As a control, it contains 49 muscle samples at different time-points of differentiation: 6 myoblasts, 5 pediatric skeletal muscle, 15 adult-young skeletal muscle, 15 adult-old skeletal muscle and 8 hSKMCs. For this analysis, we used 311 samples of tumor and healthy tissues. Gene expression data normalization was performed using RMA algorithm included in the oligo R-package (R/Bioconductor). Quality control was done using oligo and limma R-packages (R/Bioconductor).

### RNA-seq preparation and bioinformatic analysis

RNA-seq libraries were prepared from 0.5–1 μg of high-quality total RNA extracted from samples using the Illumina Stranded Total RNA Prep kit (Illumina), according to the manufacturer’s instructions. The RNA-seq samples were mapped against the hg38 human genome assembly using TopHat^63^ with the option –g 1 to discard those reads that could not be uniquely mapped in just one region. Genome-wide profiles for genome viewer visualization were generated with SeqCode^64^. DESeq2^53^ was run to quantify the expression of every annotated transcript using the RefSeq catalog of exons and to identify each set of differentially expressed genes. Reports of functional enrichments of GO and other genomic libraries were generated using the EnrichR tool^46^ and the clusterProfiler R package^65^. GSEA of the pre-ranked lists of genes by DESeq2 stat value was performed with the GSEA software^66^. Heatmaps were performed with heapmap2 R package. PCAplots were generated with the factoextra and plot3D R packages.

### Immunohistochemistry

Immunohistochemical analyses were performed following standard techniques. The primary antibodies and dilutions are listed (see Supplementary Table S6). Tumors were fixed in formalin and embedded in paraffin for subsequent processing. Next, sections were deparaffinized, rehydrated, and heated with Epitope Retrieval Solution (pH 6.0) (Novocastra Laboratories). Reactions were developed with Novolink Polymer Detection System (Novocastra Laboratories). Immunoreactivity was visualized by diaminobenzidine, and nuclei were counterstained with hematoxylin. Tissue was then dehydrated with alcohol, permeated with xylene, and mounted with Permount organic mounting solution (Thermo Fisher Scientific). Nanozoomer 2.0 (RRID:SCR_021658) was used to scan selected tumors for digital image processing. Images were evaluated by a pathologist to select regions of interest and analyzed with the Dotslide Microscope and Olympia Software (Olympus). Similar regions of every sample were selected from every section.

### Chromatin immunoprecipitation followed by quantitative polymerase chain reaction

Cells were fixed with 1% of methanol-free formaldehyde (Thermo Fisher Scientific) at room temperature for 10 min, and the cross-linking reaction was stopped by adding 500 μl glycine (1.25 M). Cells were resuspended in lysis buffer [0.1% SDS, 0.15 M NaCl, 1% Triton X-100, 1 mM EDTA, 20 mM tris (pH 8), and protease inhibitors (1 mg/ml)] and sonicated with Bioruptor Pico (Diagenode) for 30-40 cycles until chromatin was sheared to an average fragment length of 200 bp. After centrifugation, a small fraction of eluted chromatin was measured with Qubit. Starting with 30 μg of sample, immunoprecipitation for each antibody was performed overnight; the primary antibodies are listed in Supplementary Table S6. 50 μl of Dynabeads Protein A (Invitrogen) was then added and incubated for 2 h at 4°C under rotation. Immunoprecipitates were washed once with TSE I [0.1% SDS, 1% Triton X-100, 2 mM EDTA, 20 mM tris-HCl (pH 8), and 150 mM NaCl], TSE II [0.1% SDS, 1% Triton X-100, 2 mM EDTA, 20 mM tris-HCl (pH 8), and 500 mM NaCl], and TSE III [0.25 M LiCl, 1% Nonidet P-40, 1% deoxycholate, 1 mM EDTA, and 10 mM tris-HCl (pH 8)] and then twice with tris-EDTA buffer. DNA captured by the beads was eluted by adding 120 μL of a solution containing 1% SDS, 0.1 M NaHCO3 and decrosslinked at 65°C for 3 hours with gentle shaking. Genomic DNA fragments were purified with QIAquick PCR Purification kit (Qiagen) and eluted in 50-100 μL of Tris-EDTA buffer. Differences in DNA content were determined by qPCR using the QuantStudio 6 Flex (Applied Biosystems) and SYBR Green master mix (Thermo Fisher Scientific).

Each immunoprecipitation was done in triplicate, and PCR assays were performed using fixed amounts of input and immunoprecipitated DNA. For every amplicon, standard curves to calculate efficiency and melting curves to confirm single amplicons were obtained. The reported data represent real-time PCR values normalized to input DNA and are expressed as percentage (%) of bound/input signal.

### ChIP-seq preparation and bioinformatic analysis

Libraries were prepared using the NEBNext Ultra DNA Library Prep from Illumina according to the manufacturer’s protocol. Briefly, 5 ng of input and ChIP-enriched DNA were subjected to end repair and addition of “A” bases to 3′ ends, ligation of adapters, and USER excision. All purification steps were performed using AgenCourt AMPure XP beads (Qiagen). Library amplification was performed by PCR using NEBNext multiplex oligonucleotides for Illumina. Final libraries were analyzed using Agilent high sensitivity ChIP to estimate the quantity and distribution before amplification with Illumina’s cBot. Libraries were loaded onto the flow cell sequencer 1 × 50 on Illumina’s HiSeq 2500. ChIP-seq samples were mapped against the hg38 human genome assembly using BowTie2^67^ and the SAMtools^68^ were used to discard those reads that could not be uniquely mapped to just one region. Model-based analysis of ChIP-seq (MACS) was run individually on each replicate with the default parameters but with the shift size adjusted to 100 bp to perform the peak calling against the corresponding control sample ChIP^69^. DiffBind was initially ran over the peaks reported by MACS for each pair of replicates of the same experiment to generate a consensus set of peaks^41^. Next, DiffBind was run again over each pair of replicates of the same experiment, samples and inputs, to find the peaks from the consensus set that were significantly enriched in both replicates in comparison to the corresponding controls (categories, DBA_CONDITION; block, DBA_REPLICATE; and method, DBA_DESEQ2_BLOCK). SeqCode^64^ was used to produce the genome-wide profiles, identify the lists of target genes, the genomic composition of each set of peaks, the heatmaps of occupancy at each cell line and the quantification of signal strength across experiments. The UCSC genome browser^70^ was used to generate screenshots of the genomic regions. MEME^71^ was used to perform the motif analysis of each collection of peaks. We perform the term enrichment analysis of target genes with the libraries of Enrichr^45,46^. GSEA analyses were executed with the GSEA tool^66^.

### Mouse xenograft models

In vivo studies were performed after the approval of the Institutional Animal Research Ethics Committee (project number 505/24). Athymic Foxn1nu nude mice (Envigo) were injected subcutaneously with 0.5 × 10^6^ cells for RH30 shPLKO, shUSP7#seq1 and shUSP7seq#2. Cells were resuspended in 200 μl of Matrigel (Becton Dickinson) 1:1 with phosphate buffered saline and injected into both flanks (5 mice, n = 10 for seq#1 and 6 mice, n = 12 for seq#2). Antitumoral effect of P22077 was studied in Athymic-nude Foxn1nu injecting subcutaneously 1 × 10^6^ RH30 cells into both flanks. When tumor volume arrived at 150-300 mm3, 30 mice were randomly separated into three groups (10 mice, n = 20 for each group): vehicle, 10 mg/kg and 20 mg/kg of P22077. We used two different doses of P22077 to evaluate dose-dependent effect. P22077 was diluted in 2.5% DMSO, 30% PEG300 and 2% Tween-80 and sonicated before intraperitoneally administration 6 days per week for 3 weeks. Toxicity during the experiment was evaluated following standard well-fare animal condition and weight. In both experiments response was evaluated by reduction in tumor size. Tumor growth was monitored three times a week by measuring tumor volume with a digital caliper. Mice were euthanized when tumors reached a size of 2000 mm^3^ and survival curves were calculated using the Kaplan-Meier method with a log-rank test. At the end of the experiment, tumors were excised; half of each specimen was frozen in liquid nitrogen for RNA extraction, and the other was fixed in 10% formalin for immunohistochemistry experiments.

### Statistical analysis

To compare quantitative variables among more than two groups we used the one-way ANOVA with Dunnett’s post-hoc test, which compares each condition with the control condition. To estimate survival curves in the data extracted from public GEO databases, we used the Kaplan-Meier method. To estimate optimal thresholds for the expression of the genes of interest^6^, we used the method of Contal-O’Quigley^72^. This allowed us to determine two subgroups of low and high expression for each gene. Survival curves were compared with the logrank test. All p-values under 0.05 were considered statistically significant. All statistical analyses were performed with GraphPad Prism 10 and R version 4.5.1^73^.

## Data availability

ChIP-seq and RNA-seq raw data and processed files (profiles and peaks on hg38) are deposited in the NCBI GEO entry GSE324327.

## ACKNOWLEDGEMENTS

Our studies were supported by grants from AGAUR and the Spanish government through the Instituto de Salud Carlos III (FI21/00047 to Táboas P.). We are grateful to the Band of Parents at Hospital Sant Joan de Déu for supporting the overall research activities of the Pediatric Cancer Group (Institut de Recerca Sant Joan de Déu). The **L.D.C**. laboratory was supported by Fundació La Caixa Health Research Grant (CI24-10473); and Worldwide Cancer Research (2025_038). The laboratory of **T.G.P.G**. and **F.C.A**. acknowledges funding by the Cancer Grand Challenges partnership (PROTECT team) supported funded by Cancer Research UK, the National Cancer Institute, the Scientific Foundation of the Spanish Association Against Cancer And KiKa (Children Cancer Free Foundation), and is co-funded by the European Union (ERC, CANCER-HARAKIRI, 101122595). Views and opinions expressed are however those of the authors only and do not necessarily reflect those of the European Union or the European Research Council. Neither the European Union nor the granting authority can be held responsible for them **T.G.P.G** acknowledges support by scholarships of the Heinrich F. C. Behr foundation and the German Academic Scholarship Foundation.

