## Supplementary Figures for "USP7 sustains PAX3::FOXO1 enhancer reprogramming and represents a therapeutic vulnerability in Rhabdomyosarcoma"

**Supplementary Figure 1. USP7 is overexpressed in FP-RMS and has prognostic value. A.** USP7 mRNA expression in cell lines obtained from the Cell Line Protein Atlas. **B.** RING1B mRNA expression in cell lines obtained from the Cell Line Protein Atlas. **C.** Immunohistochemistry (IHC) of RING1B in three TMA representative samples from Hospital Sant Joan de Déu. Images were acquired at 5X (top) and 20X (bottom) magnifications using a Leica CTR5000 microscope. Table shows RING1B levels score of IHC samples. **D.** Correlation heatmap comparing the 101 RMS samples from E-TABM-1202 database. **E.** Survival of RMS patients based on the FP and FN status ($p$ = 0.00011). **F.** Survival of RMS patients comparing high ($\geq 4.28$, n = 29) versus low ($< 4.28$, n = 72) levels of RING1B (p = 0.13).

**Supplementary Figure 2. USP7 and RING1B colocalize with PAX3::FOXO1 at active enhancers. A.** Volcano plots of the significant peaks identified by DiffBind for PAX3::FOXO1 (P3F1), USP7 and RING1B for each pair of replicates. Significant peaks (pink dots) were selected by $p < 0.05$; FDR $< 10^{-4}$. Non-significant peaks are represented in blue dots. **B.** Boxplots depicting the ChIP-seq signal strength of the PAX3::FOXO1 antibody in the FP-RMS (RH30, RH4 and ARMS-006) and the FN-RMS (RD) cell lines at each set of PAX3::FOXO1 peaks identified in the corresponding FP-RMS cell line. **C.** Bar plot depicting percentage of peaks over regulatory elements (active/poised enhancer and promoters) for PAX3::FOXO1 (P3F1), RING1B and USP7 (left) and for histone marks H3K4me3, H3K4me1, H3K27ac, and H3K27me3 (right, as a control) in RH4 cell line. White represents active enhancers, light grey active promoters, dark grey poised enhancers and black poised promoters. **D.** Heatmap of PAX3::FOXO1, USP7, RING1B,

H3K27ac and negative control (input) ChIP-seq signals normalized by the total number of mapped reads at the PAX3::FOXO1 peaks in RH4 and HSJD-ARMS-006 models. **E.** Bar chart representing the top ten ChEA 2022 enriched categories of genes associated with PAX3::FOXO1 (left), USP7 (middle) and RING1B (right) peaks in RH30 cells. Bar length indicates the -log10 (adjusted p value). All shown pathways have a p value < 0.05. Y axis shows the top 10 enriched pathways provided by EnrichR tool[45,46]. **F.** Pie chart showing genomic distribution of KDM2B TOP-5000 peaks in RH30, RH4 and HSJD-ARMS-006. The three areas are promoters (blue), intragenic (grey) and intergenic (white) regions. **G.** Venn diagram depicting the overlap of genes associated with PAX3::FOXO1, USP7, RING1B and KDM2B peaks in RH30 cells. **H.** Boxplots comparing RING1B and RING1A ChIP-seq signal strength on RH30 within PAX3::FOXO1 peaks.

**Supplementary Figure 3. USP7 downregulation reduces tumorigenic capacity of FP-RMS cells. A.** Western blot showing USP7 levels upon shPLKO, shUSP7 #1 and #2 in RH30, RH4 and HSJD-ARMS-006. Cells were selected with puromycin for 10 days and tubulin was used as a loading control. **B.** Western blot showing protein levels of PRC1.1 subunits KDM2B, USP7, RING1A, RING1B and RYBP in RING1B KO (left) and USP7 KD (right) in RH30. Tubulin was used as a loading control. **C.** UCSC ChIP-seq signal tracks for PAX3::FOXO1 (P3F1, red), RING1B (blue), USP7 (green), KDM2B (orange) and H3K27ac (pink) in RH30, RH4 and HSJD-ARMS-006 for KDM2B genomic region, normalized by the total number of mapped reads. **D.** RT-qPCR determination of mRNA relative expression of PRC1.1 subunits genes (*RING1A, RING1B,* and *USP7), PAX3::FOXO1,* CRC genes *(MYOD1, MYOG, MYCN),* and oncogene targets *(ALK, FGFR4, FOXF1, IGF1R* and *JARID2)* in sgRING1B #1 and #2 compared to control in RH30 cells. TBP was used as a housekeeping gene for normalization. Significant p values were shown in the graph. **E.** RT-qPCR determination of mRNA relative expression of *MYH1* and *MYF5* in shUSP7 #1 and #2 compared to control in RH30 cells. Significant p values were shown in the graph. **F.** RT-qPCR determination of mRNA relative expression of *MYH1* and *MYF5* in sgRING1B #1 and #2 compared to control in RH30 cells. Significant p values were shown in the graph. **G.** Cell counts of RH30 wild type cells compared to sgRING1B #1 and sgRING1B #2. **H.** Representative replicates of colony formation capacity in RH30 wild type versus sgRING1B #1 and #2 ($p$ = 0.0007 and 0.0013, respectively). Quantification and p values of all replicates are shown.

**Supplementary Figure 4. USP7 inhibitor P22077 reduces FP-RMS tumor growth *in vivo*. A.** Western blot showing H2BK120ub levels in RH30 and RH28 cells treated with P22077 at their IC50 concentration. H3 was used as a loading control. **B.** Western blot showing increasing levels of cell death marker cPARP at increasing doses of P22077 (ranging from 2 to 10 μM) in RH30 cells. Tubulin was used as a loading control. **C.** Western blot showing cPARP and USP7 levels after U7D-1 treatment at 0.5 μM from 2 to 24 hours. **D.** Western blot showing KDM2B, USP7, RING1A, RING1B and RYBP levels upon P22077 treatment in RMS cells RH4, RH30 and RH28. Tubulin was used as a loading control. **E.** Western blot showing histone protein levels of H3K27me3, H3K27ac, H2AK119ub and γH2AX upon P22077 treatment in FP-RMS cell lines RH4, RH30 and RH28. H3 was used as a loading control. **F.** Western blot showing protein levels of USP7 and p21 (left) and PAX3::FOXO1 (right) in RH30 cells treated with P22077 for 24 h. Tubulin was used as a loading control. **G.** Percentage change in weight across the experimental groups. Weight changes were calculated relative to the weight of each animal the day of the first dose of the treatment.

**Supplementary Figure 5. USP7 inhibition reduces the expression of PAX3::FOXO1 targets in FP-RMS. A.** Principal Component Analysis (PCA) of DMSO (blue) and P22077 (orange) conditions in RH4, RH30 and RH28 from RNA-seq experiments. **B.** Bar chart representing the number of genes upregulated and downregulated upon P22077 treatment in RH30, RH4 and RH28 cells. **C.** Bubble chart depicting the top 12 enriched gene ontology (GO) Biological Processes of upregulated genes upon USP7 inhibition in RH28 (left) and RH4 (right) cells. Categories were ranked by the fraction of the gene set belonging to this function. Circle size represents the number of genes on the list belonging to this function. Circle color denotes the adjusted P value of each enrichment. **D.** Bar chart representing the top ten ChEA 2022 enriched categories of downregulated genes by RNA-seq in RH28 upon USP7 inhibition with P22077. **E.** RT-qPCR determination of mRNA relative expression of PRC1.1 subunits genes, *PAX3::FOXO1,* CRC genes and oncogene targets in DMSO versus P22077 conditions in HSJD-ARMS-006. TBP was used as a housekeeping gene for normalization. Significant p values are shown in the graph. **F.** RT-qPCR determination of mRNA relative expression of PRC1.1 subunits genes, *PAX3::FOXO1,* CRC genes and oncogene targets in control versus 0.5 µM U7D-1 conditions. TBP was used as a housekeeping gene for normalization. Significant p values were shown in the graph.

**Supplementary Figure 6. USP7 inhibition induces myogenic differentiation in FP-RMS. A.** Heatmap showing the expression pattern of genes from the Molecular Signatures Databases (MSigDB, gene set Hallmarks_Myogenesis) in RH28 (left) and RH4 (right). Gene expression values are displayed as row-scaled normalized expression (z-scores), where red indicates higher expression and blue indicates lower expression relative to the mean across samples. Columns represent individual biological replicates and rows represent genes. **B.** Western blot showing MYH1 levels upon P22077 treatment in HSJD-ARMS-006 cells treated at their IC50 of P22077. GAPDH was used as a loading control. **C.** RT-qPCR determination of mRNA relative expression of *MYH8, MEOX1* and *TNNT3* in control versus P22077 conditions. TBP was used as a housekeeping gene for normalization. **D.** RT-qPCR determination of mRNA relative expression of *MYH1* and *MYF5* in control versus 0.5 μM U7D-1 conditions. TBP was used as a housekeeping gene for normalization.

### Supplementary Figure 1

A

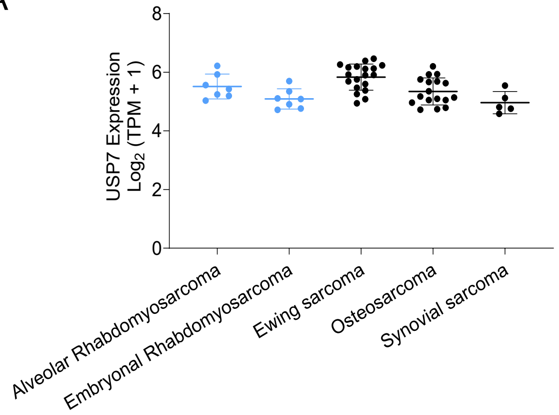


B

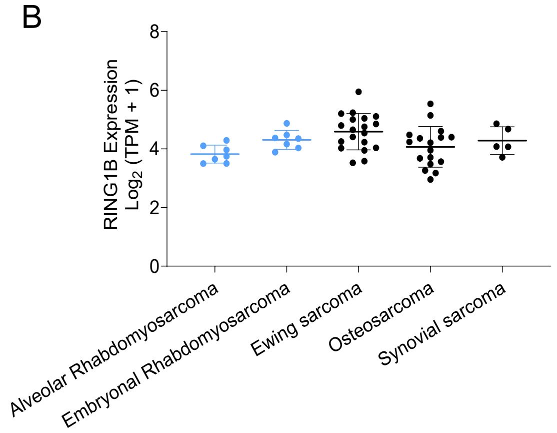


C

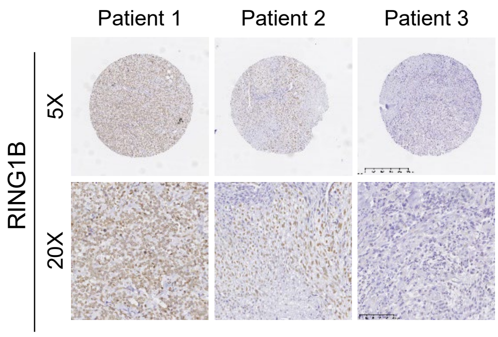


| RING1B levels | Number of patients |
|---|---|
| No levels | 3 |
| Low levels | 7 |
| Medium levels | 10 |
| High levels | 3 |

D

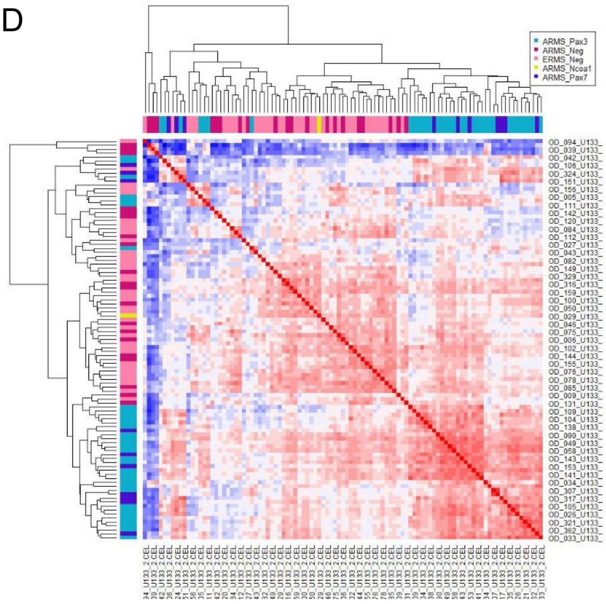


E

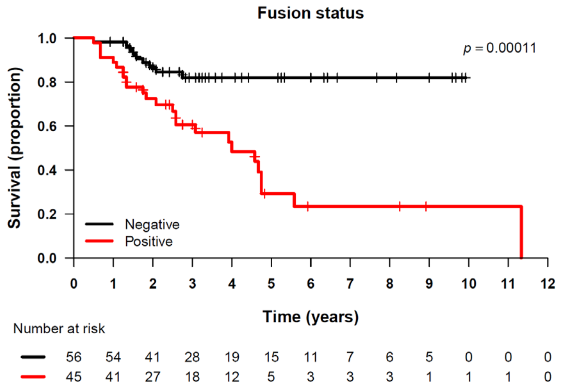


F

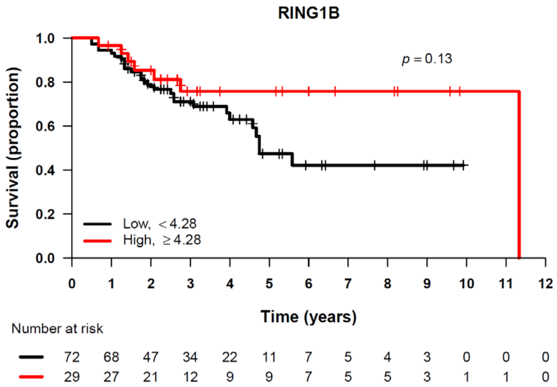

### Supplementary Figure 2

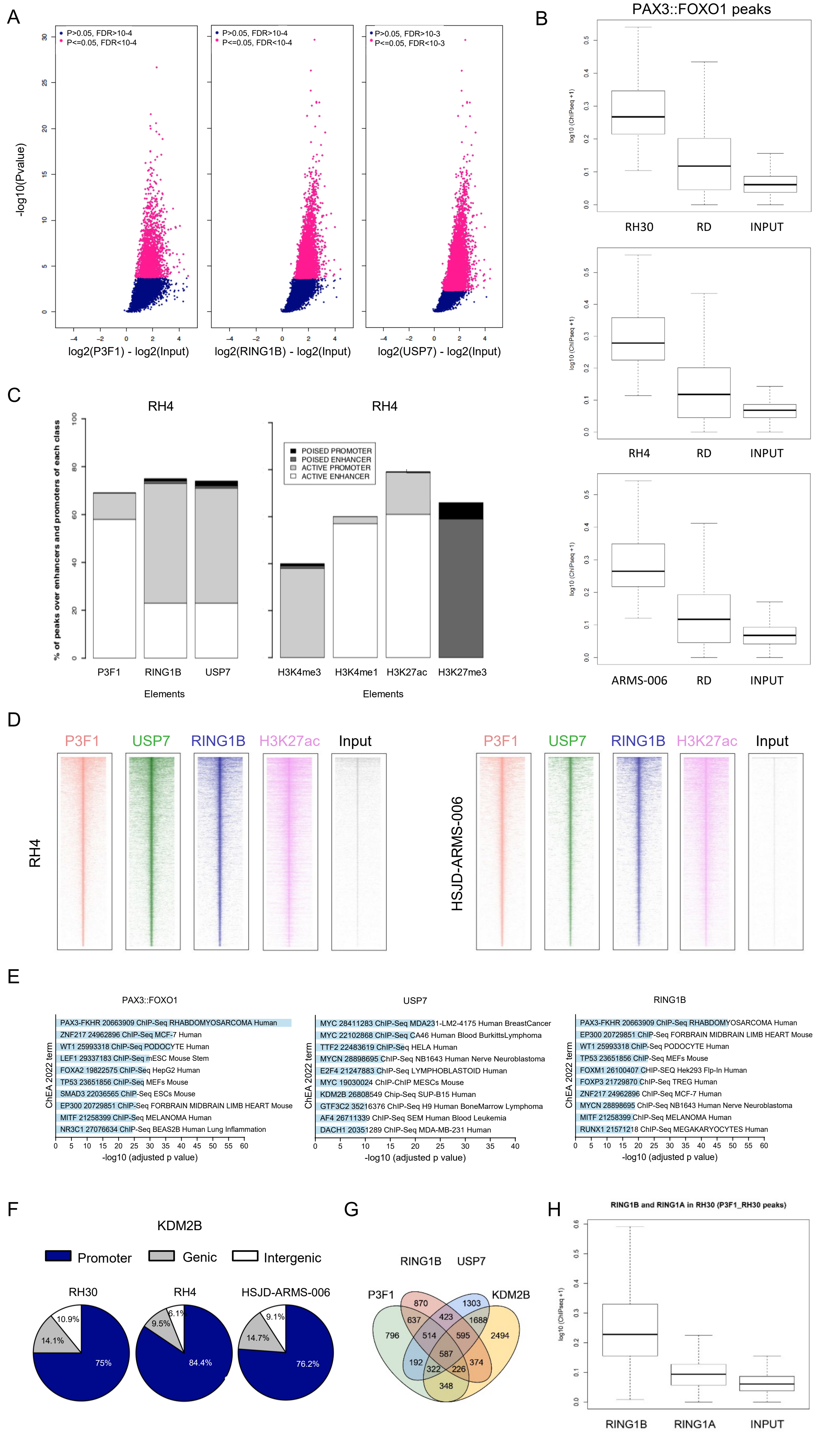

### Supplementary Figure 3

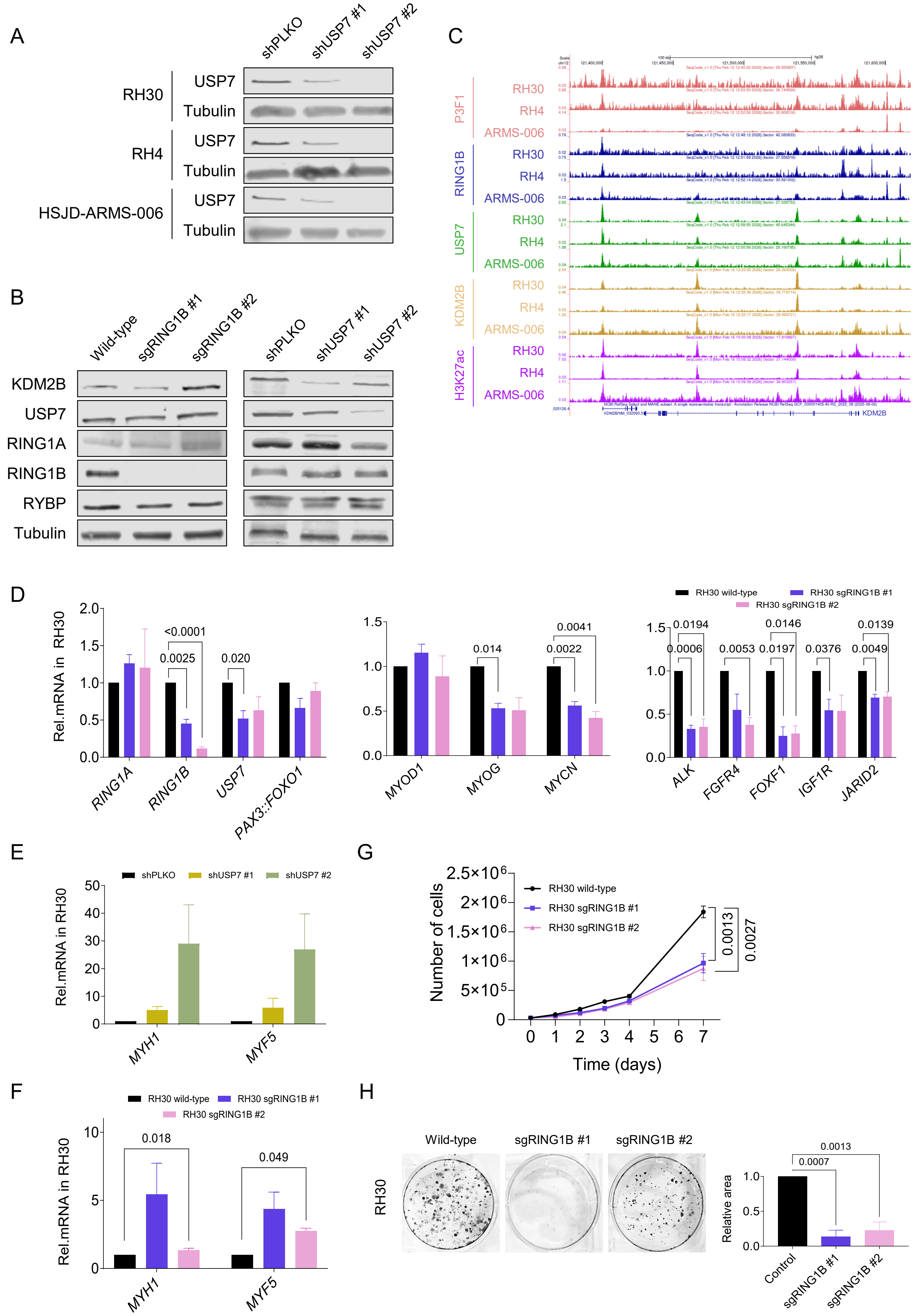

### Supplementary Figure 4

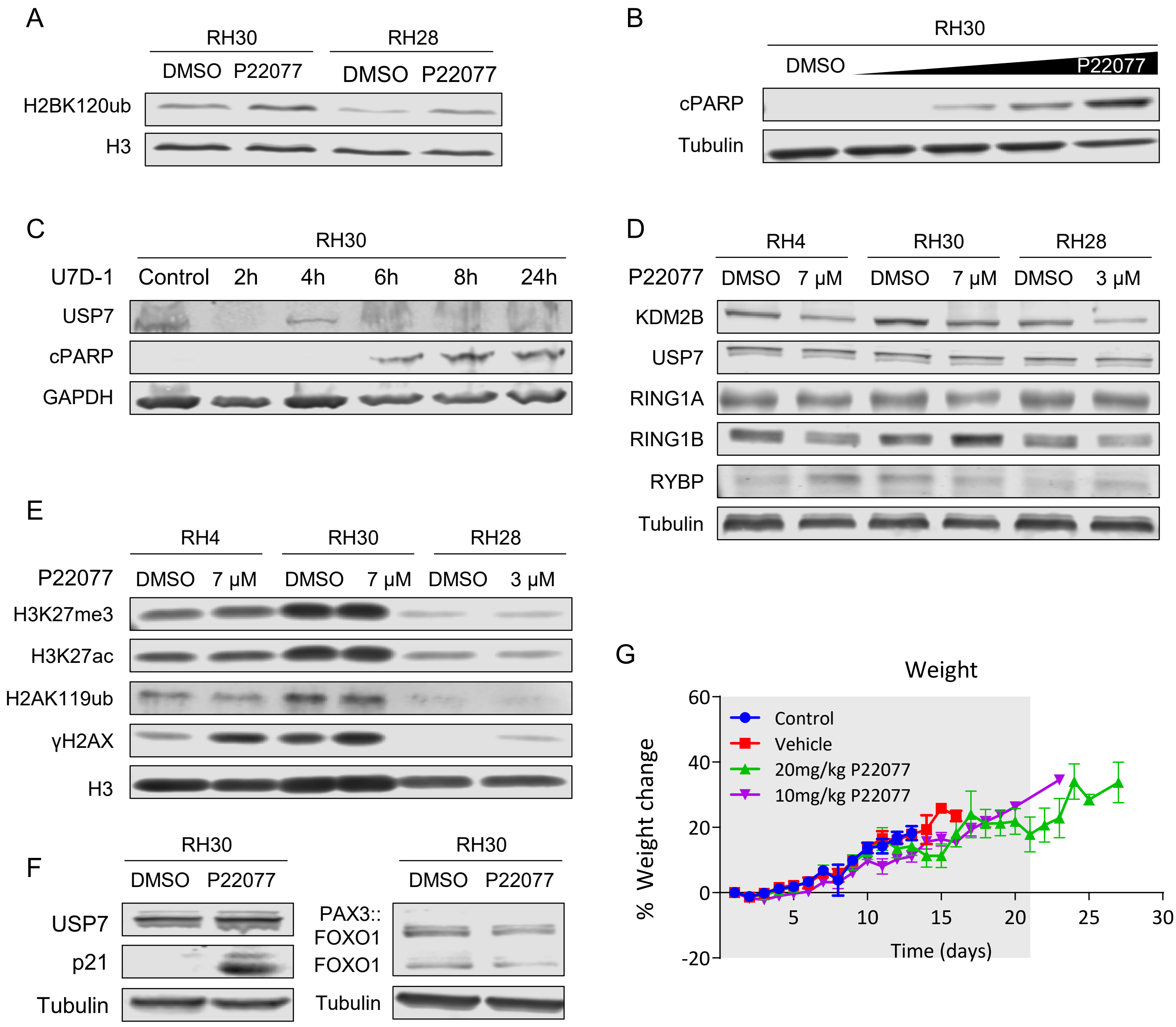

### Supplementary Figure 5

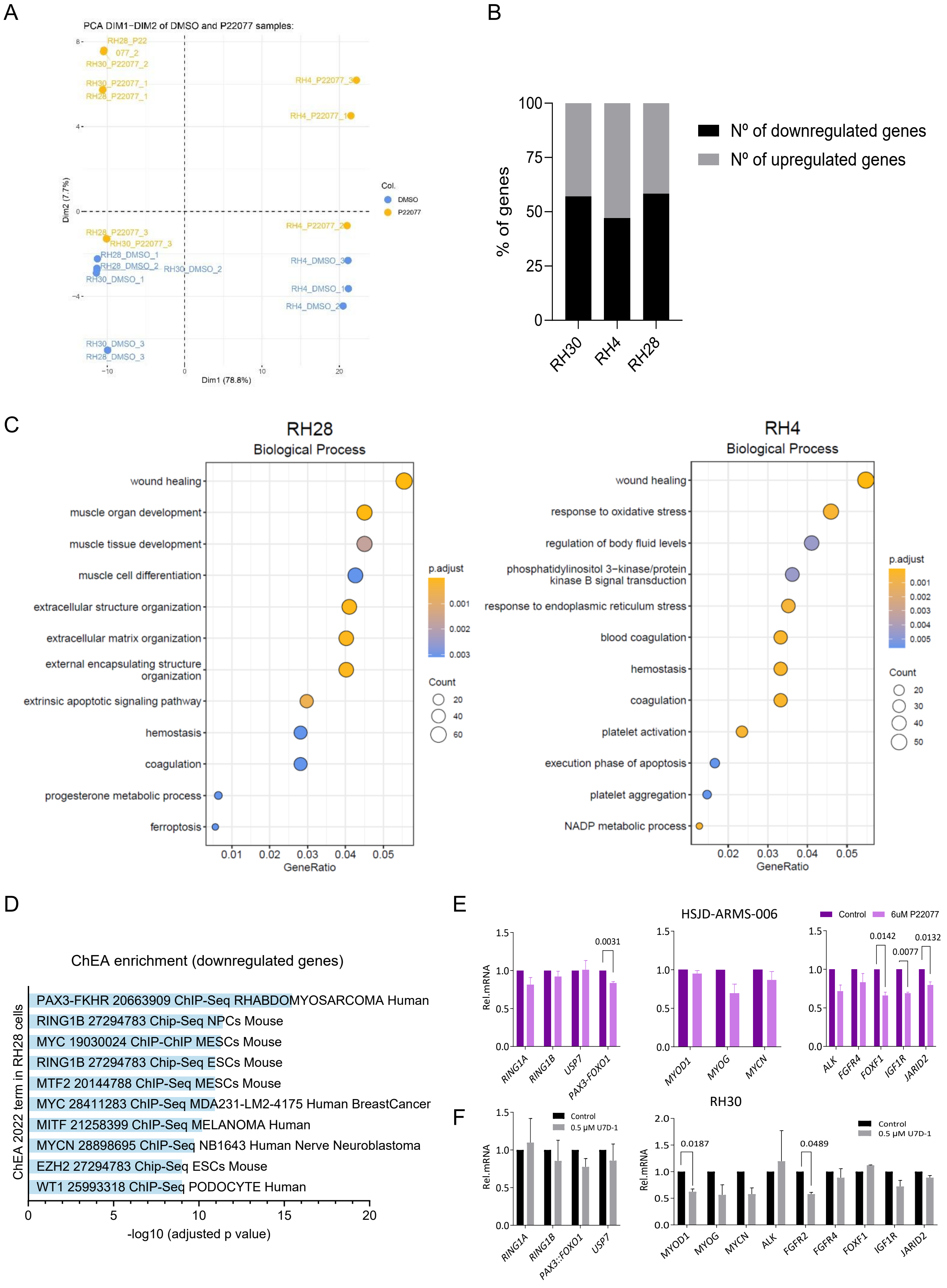

### Supplementary Figure 6

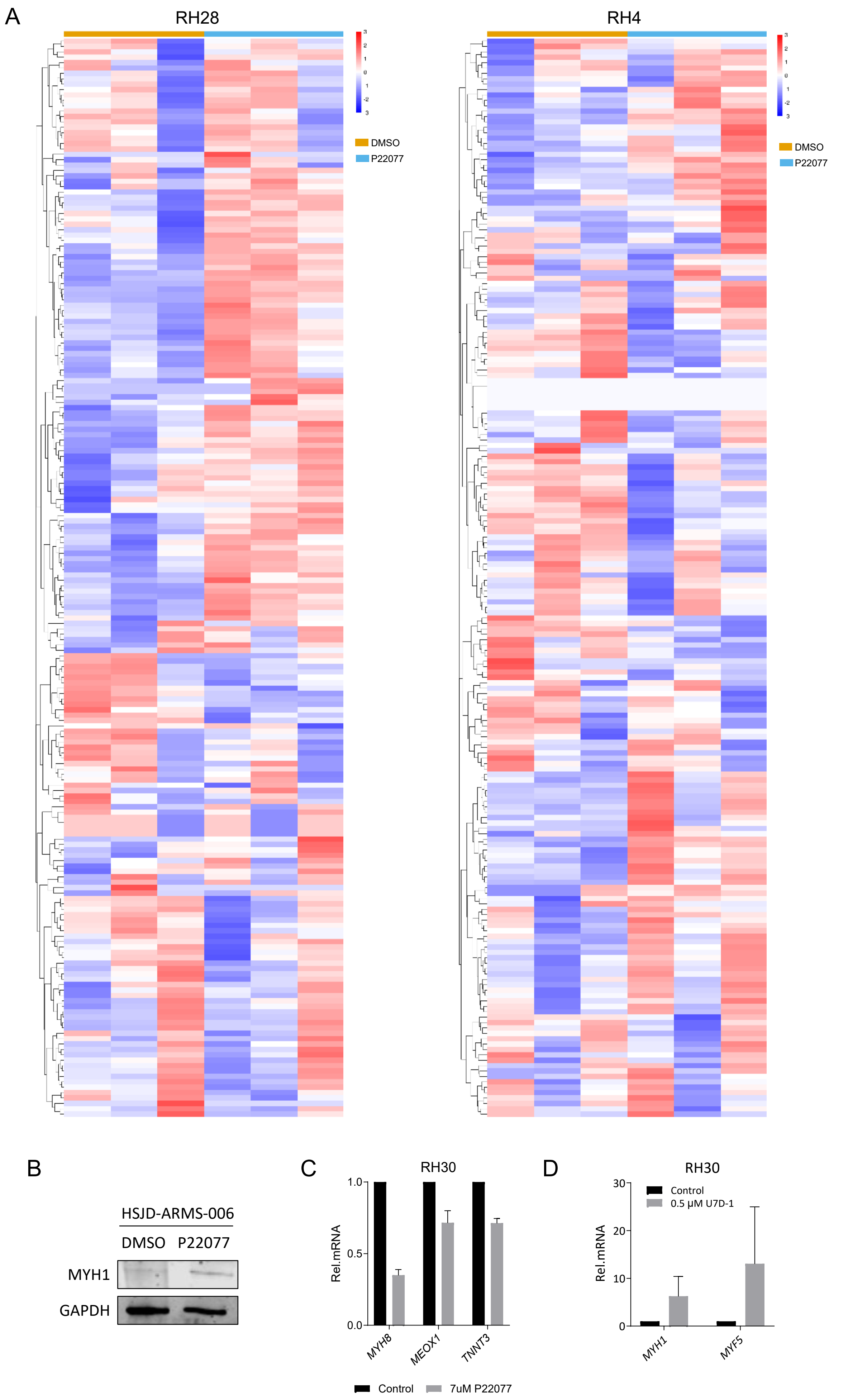
